# Gcn5 and mTORC1 mediated regulation of autophagy controls *Drosophila* blood cell homeostasis

**DOI:** 10.1101/2023.07.04.547637

**Authors:** Arjun Aji Reeja, Mincy Kunjumon, Suraj Math, Laxmi Kata, Rohan Jayant Khadilkar

## Abstract

Blood progenitors are regulated by a variety of systemic and nutritional cues from their environment. In the *Drosophila* lymph gland (LG), the Posterior Signalling Center (PSC) acts as a stem cell niche striking a balance between progenitors and differentiated blood cells. Autophagy is a vital cellular process that maintains homeostasis by removing unnecessary or dysfunctional cell components through autophagic degradation and recycling. Here, using genetic perturbation analysis, we show that autophagy plays a critical role in regulating LG blood cell homeostasis. General control non-derepressible 5 (Gcn5), a histone acetyltransferase is expressed in the primary LG lobe and modulation of Gcn5 levels perturbs LG homeostasis. Our results show that hemocyte-specific Gcn5 modulation controls autophagic flux in the hemocytes. Furthermore, we show that modulation of mTORC1 activity can perturb hematopoiesis. Our results indicate that organismal Gcn5 levels respond to dietary shifts and are modulated by mTORC1 signaling. Chemical intervention shows that mTORC1 over-rides the effect exerted by Gcn5 in regulating LG hematopoiesis. Taken together, our findings demonstrate that Gcn5 and mTORC1 regulates autophagy to maintain blood cell homeostasis in *Drosophila*.

## Introduction

*Drosophila* hematopoiesis occurs in two different waves: the first wave occurs in the embryonic stages in the procephalic mesoderm giving rise to both circulating and sessile pool of hemocytes. While the second wave of definitive hematopoiesis begins in the dorsal mesoderm, giving rise to the lymph gland (LG), the larval hematopoietic organ ^1,2^. The larval LG can be divided into functionally distinct regions: the Posterior Signaling Center (PSC), which acts as the stem cell niche producing signals that are not only required for maintaining the prohemocytes but also for priming them towards differentiation. The Medullary Zone (MZ) is made up of prohemocytes (hematopoietic progenitors) that are capable of differentiating into three differentiated blood cell types namely plasmatocytes, crystal cells, and lamellocytes. The Cortical Zone (CZ) houses the differentiated blood cell population ^3^. Single cell sequencing efforts have identified heterogeneity amongst the developing hemocytes in the LG and have discovered previously unreported hemocyte types like adipohemocytes, stem-like prohemocytes, intermediate prohemocytes, early lamellocytes, suggesting that there is a lot of heterogeneity in the cell types present within the LG for example the subsets identified within the progenitors: core, distal and intermediate ^4,5^. These recent single cell sequencing based studies show that the LG is a complex organ with multiple hemocyte types and consists of an intricate array of signals that regulate them ^4,6^.

The cellular fate of prohemocytes in the medullary zone is intricately controlled by a set of extrinsic and intrinsic signals. Multiple signaling pathways like the JAK/STAT, Hedgehog, Wingless, Dpp have been reported to play an important role in prohemocyte maintenance ^7–15^. Stem cell and tissue homeostasis is closely linked to the nutritional status of the organism. There is increasing evidence indicating the involvement of nutrient signals in stem/progenitor cell fate ^16–19^. *Drosophila* hematopoietic progenitors are capable of sensing systemic and nutritional cues. Starvation has been shown to alter the homeostatic balance in the LG mimicking an inflammatory response like scenario causing excessive differentiation of blood cells ^20^. Also, perturbation of key nutrient sensing pathways like the IIS/Tor signaling has been shown to affect niche numbers as well as progenitor differentiation ^21^. More specifically TORC1 has been shown to play an important role during *Drosophila* hematopoiesis. TORC1 activity has been shown to promote the proliferation and expansion of progenitor pools^22^. Knocking down TOR signaling components in the *Drosophila* hematopoietic niche has shown to cause a cell autonomous reduction in the niche numbers. Activation of Tor signaling on the other hand results in excess progenitor differentiation. This shows that a fine tuning of the nutrient sensing pathways is important for progenitor maintenance ^23^. Autophagy a cell intrinsic process is also known to be responsive to nutritional status and stress. Autophagy ensures cellular homeostasis by removing intracellular waste material, replaces old and damaged organelles and sustains cell survival during nutrient deprived conditions ^24,25^. mTOR (molecular target of rapamycin) is an important regulator of autophagy. mTORC1 inhibits autophagy in nutrient replete conditions by targeting either initiation, expansion or termination of autophagy ^26^. Several studies indicate that autophagy is essential for the maintenance of hematopoietic stem cells ^27–29^. There have been several reports that investigate the role of key autophagy genes in hematopoiesis and show that mice lacking functional autophagy genes like Atg7, Atg12 or FIP200 lead to an exhaustion of the HSC pool especially long-term HSCs thus showing that autophagy is critical for HSC maintenance ^28,30,31^. Furthermore, HSCs lacking Atg12 displayed phenotypes similar to aged HSCs viz. increased mitochondrial content, heightened metabolic activity, myeloid differentiation bias ^27^. Similarly, autophagy genes are shown to affect *Drosophila* hematopoiesis, it has been shown that Atg6 mutants show an enlarged LG and possess elevated blood cell numbers ^32^. Perturbing autophagy by knocking down Atg1 affected hemocyte spreading and recruitment to wound site^33^. Inhibiting Atg2, induced melanotic nodule formation, impaired the ability to respond to bacterial infection by altering the expression of AMP (Anti -Microbial peptides) genes as well as disrupting phagocytosis ^34^.Mis-regulation of the autophagic process can lead to leukemogenesis^35^. Though the role of autophagy has been explored in mammalian hematopoiesis, its role and mechanisms it controls during *Drosophila* hematopoiesis is unexplored. There are various nutrient sensors that can sense systemic nutrient signals and regulate cellular processes like autophagy. In this study, we report that General control non-depressible 5 (GCN5), a histone acetyltransferase, regulates the process of autophagy during hematopoiesis through its non-histone acetylation target, TFEB.

General control non-depressible 5 (GCN5) is the first nuclear histone acetyltransferase (HAT) to be identified with function in transcriptional regulation by serving as a key subunit in transcriptional coactivator complexes like SAGA (Spt-Ada-Gcn5-acetyltransferase) and ATAC (Ada-2A-containing) ^36–38^. The transfer of acetyl group to the lysine residues of histones mediates HAT activity^39^ . The primary acetylation site of Gcn5 is lysine 14 of Histone 3 (H3K14) whereas other lysine residues of Histone 3 like H3K9, H3K18, H3K23, H3K27 are also found to be acetylated by Gcn5^40^ . Recent studies reported the additional role of Gcn5 in histone succinylation as well as crotonylation^41–43^. Gcn5 has important functions in the extension of lifespan^44^, metabolism ^45^, cellular differentiation^46^, cell proliferation and growth^47^, neuronal apoptosis^48^ and DNA damage response ^49^. GCN5 acts as an oncoprotein in cancer and its inhibition can rescue cancer phenotypes in various cancers namely hepatocellular carcinoma, glioma, non-small cell lung carcinoma^50–52^ . Gcn5 is over-expressed in Acute Myeloid Leukemia (AML) and is needed for the survival of AML cell lines ^53^. Aberrant acetylation of H3K9 by GCN5 leads to all-trans Retinoic acid (ATRA) resistance in non-APL AML, thereby preventing the maturation of myeloid blasts and maintaining them in a leukemic state^54^ . GCN5 plays a role in drug resistance in the leukemic early drug-resistant population (EDRP) where it interacts with ATM thereby recruiting ATM to the DNA damage site ^55^. With an evolutionary well conserved structure and function, the significance of Gcn5 in different cellular processes can be studied using a variety of model organisms^42^ . *Drosophila* has only one homolog of Gcn5 (dGCN5), hence making it an efficient model to study the functions of Gcn5 ^56^. In *Drosophila*, loss of Gcn5 function affects developmental processes like oogenesis, metamorphosis^57^ . Gcn5 plays an important role in germline stem cell maintenance^58^ . Other than the developmental processes, Gcn5 was shown to regulate a cellular process like autophagy by targeting TFEB, a non-histone acetylation target of Gcn5 that activates genes responsible for autophagosomal and lysosome biogenesis ^59^. mTORC1 regulates autophagy not only through the initiator kinase, Ulk1 arm but also through inhibitory phosphorylation of TFEB ^60^. In the context of hematopoiesis, Gcn5 is necessary for T-cell development and function, and deletion of Gcn5 could affect the development of immune cells in vertebrates^46^ . Gcn5 expression increases during the differentiation phase of human CD34 cells indicating an important role of Gcn5 in erythroid differentiation^61^ . Although there are studies demonstrating the role of Gcn5 in leukemogenesis there is paucity of information on its role in physiological hematopoiesis. In *Drosophila*, a study reported that loss of a HAT like Gcn5 or Chameau phenocopies the differentiation defect seen upon loss of function of molecules regulating fatty acid oxidation^62^ . There are no other studies that have explored the role of Gcn5 during normal hematopoiesis in *Drosophila*.

Here, we show that Gcn5 plays an essential role in regulating blood cell homeostasis in the *Drosophila* LG. Analysis of *gcn5* mutants, cellular subset specific modulation of Gcn5 levels in the LG and by utilizing structure function approaches in the LG, we show that Gcn5 function is important for developmental hematopoiesis. Our results demonstrate that modulation of Gcn5 levels alter blood cell homeostasis via its regulation of autophagy. We show that autophagy plays a critical role in the LG by controlling blood cell differentiation. Gcn5 acts as a nutrient sensor and negatively regulates autophagy through its non-histone acetylation target, TFEB. Additionally, our findings indicate that modulation of mTORC1 activity can regulate Gcn5 levels and perturb LG hematopoiesis. mTORC1 over-rides the effect exerted by Gcn5 in regulating LG hematopoiesis. Taken together, we suggest that the Gcn5 – mTORC1 – TFEB signaling axis mediated control of autophagy is required for maintaining blood cell homeostasis in *Drosophila*.

## Results

### Gcn5 is expressed in all the cellular subsets of the primary LG lobe

Gcn5 has been previously reported to be overexpressed in acute myeloid leukemias (AML) and is required for the survival of AML cells ^53–55^. In *Drosophila*, Gcn5 has been shown to play an important role during metamorphosis and in the maintenance of germline stem cells^57,58^ . However, its role in *Drosophila* hematopoiesis is unknown. We first set out to understand the expression of Gcn5 in the *Drosophila* LG where the primary lobe is the main site of active hematopoiesis. The primary lobe consists of various developmental zones housing different cell populations with identifiable markers (Fig. 1A, B). Gcn5 expression analysis shows that Gcn5 is expressed in all cellular subpopulations of the primary LG lobe including the Posterior Signalling Center (PSC, Fig 1C-C’, E-E’’), Dome positive medullary zone (MZ) progenitors and Dome negative hemocytes in the LG (Fig 1D-D’, F-F’). Gcn5 is also expressed in the posterior LG lobes (Fig. S5E-F).

**Figure 1.**
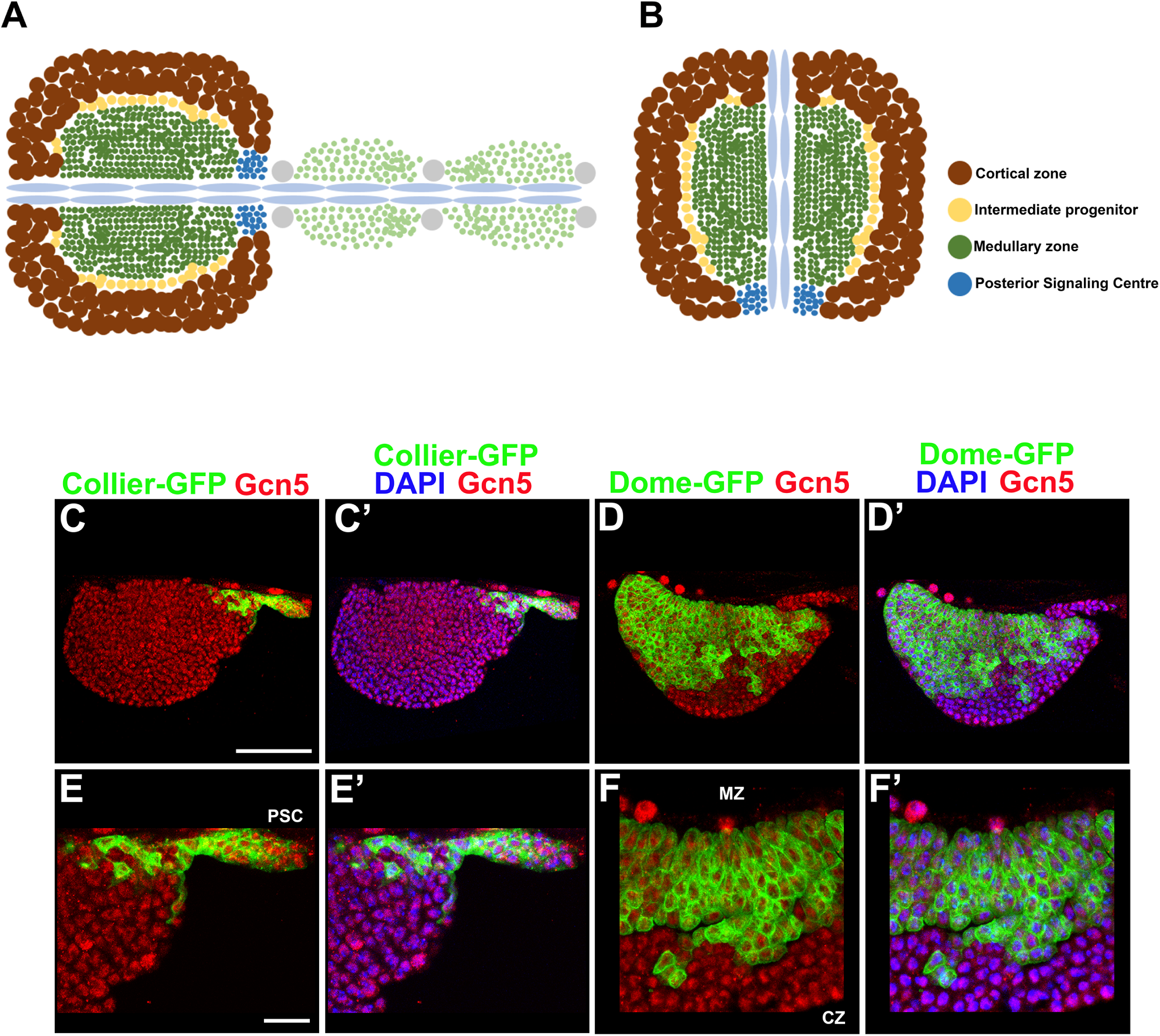
Gcn5 is expressed in all compartments of the *Drosophila* larval LG. Cartoon of the third instar larval LG, indicating the anterior primary lobes and the posterior lobes (A). The primary lobe of the LG depicting different subpopulations: the Posterior Signaling Centre (PSC: blue) functioning as the niche, Medullary zone (MZ: green) houses the progenitors, Intermediate progenitor zone (IZ: yellow) which contains the cells transitioning from progenitors to specific lineages, and the Cortical zone (CZ: brown) containing the differentiated hemocytes population – plasmatocytes and crystal cells (B). Gcn5 expression in different compartments of lymph gland (C-F’). Gcn5 (red) expression in the - Posterior Signaling Centre (PSC) using Collier- GFP (green) (C-C’, E-E’), Medullary zone prohemocytes using Dome-GFP (green) (D-D’, F-F’). Nuclei are stained with DAPI (Blue). Scale Bar: 50µm (C-D’), 30µm (E-F’).

### Whole animal *gcn5* mutants show defective blood cell homeostasis

Since Gcn5 is expressed in all the cellular subpopulations of the LG, we were interested to know if hematopoiesis gets affected in the whole animal *gcn5* mutants. We first characterized the hematopoietic phenotypes of the *gcn5* null allele, *gcn5[E333st]* and the hypomorphic allele, *gcn5[C137Y]* in heterozygous conditions. We observed that the Antennapedia positive PSC cell numbers were not affected in the *gcn5[C137Y/+]* or *gcn5[E333st/+]* heterozygous mutants compared to wildtype (Fig S1A-D). Excessive differentiation of P1 (NimRodC1) positive plasmatocytes was observed in the *gcn5[E333st/+]* mutant whereas no significant change was seen in the *gcn5[C137Y/+]* mutant compared to wildtype (Fig S1E-H). In contrast, a significant increase in Hindsight positive Crystal cell differentiation and gamma-H2Ax positive DNA damage foci were observed in both *gcn5[C137Y/+]* and *gcn5[E333st/+]* mutants, compared to wildtype (Fig S1I-P). We assessed the impact of Gcn5 perturbation on DNA damage accumulation due to its known role in regulating DNA damage repair ^49,63,64^. We then combined these alleles in homozygous and trans-heterozygous combinations. The *gcn5 [C137Y]* hypomorphic allele and the null allele *gcn5 [E333st]* null allele were combined which yielded viable homozygous larvae whose LG hematopoietic phenotypes were studied. We also generated *gcn5 [C137Y]/gcn5 [E333st]* transheterozygotes in order to study the hematopoietic phenotypes of the *gcn5* mutants. Our analysis reveals that the PSC cell numbers were reduced in the *gcn5[E333st/C137Y]* transheterozygous mutant as well as the *gcn5[E333st/ E333st]* mutant whereas no change was observed in the *gcn5[C137Y/C137Y]* homozygous mutant, compared to wildtype (Fig 2A-E). Plasmatocyte differentiation significantly increased in the *gcn5[C137Y/C137Y]* and *gcn5[E333st/ E333st]* homozygous mutants, whereas no significant change was observed in *gcn5[E333st/C137Y]* transheterozygous mutant compared to wildtype (Fig 2F-J). A significant increase in crystal cell differentiation and DNA damage was observed among all the three mutants *gcn5[C137Y/C137Y]*, *gcn5[E333st/C137Y]* and *gcn5[E333st/ E333st]* respectively, compared to wildtype (Fig 2K-T). Since these are all whole animal mutants of Gcn5, there could be a number of systemic signals that could contribute to these changes in hematopoiesis as the LG is very sensitive to systemic signals and responds rapidly. Further characterization of systemic signals and signaling mechanisms that are causing these aberrations in hematopoiesis needs to be done,

**Figure 2.**
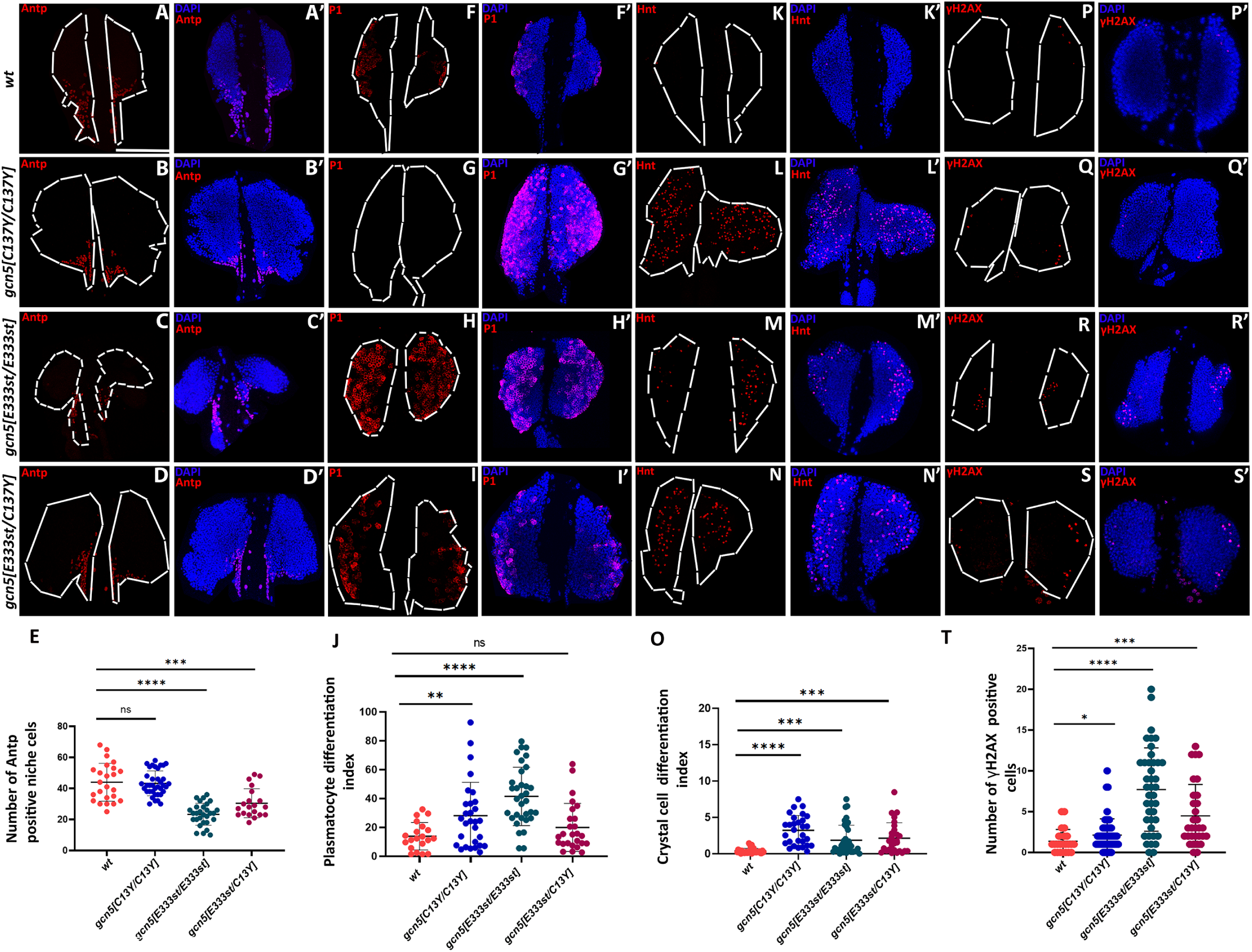
Whole animal *gcn5* mutants show defective LG homeostasis. Posterior Signaling Center (PSC) cell numbers marked by Antennapedia (red) in *gcn5[C137Y/C137Y]* (B-B’), *gcn5[E333st/E333st]* (C-C’) and *gcn5[E333st/C137Y]* (D-D’’) as compared to wildtype (A-A’). Graphical representation of PSC cell numbers of the *gcn5* mutants as compared to the wildtype (E), n=24 for wildtype, n=37 for *gcn5[C137Y/C137Y]*, n=29 for *gcn5[E333st/E333st]* and n=20 for *gcn5[E333st/C137Y]* for PSC cell number quantification. Plasmatocyte differentiation was marked by P1 (red) in *gcn5[C137Y/C137Y]* (G-G’), *gcn5[E333st/E333st]* (H-H’) and *gcn5[E333st/C137Y]* (I-I’) as compared to the wildtype (F-F’). Graphical representation of plasmatocyte differentiation index of the *gcn5* mutants compared to wildtype (J), n=20 for wildtype, n=29 for *gcn5[C137Y/C137Y]*, n=33 for *gcn5[E333st/E333st]* and n=26 for *gcn5[E333st/C137Y]* for quantification. Crystal cell differentiation marked by Hnt (red) in *gcn5[C137Y/C137Y]* (L-L’), *gcn5[E333st/E333st]* (M-M’) and *gcn5[E333st/C137Y]* (N-N’) as compared to wildtype (K-K’). Graphical representation of crystal cell differentiation index for *gcn5* mutants compared to wildtype (O), n=34 for wildtype, n=29 for *gcn5[C137Y/C137Y]*, n=38 for *gcn5[E333st/E333st]* and n=28 for *gcn5[E333st/C137Y]* for quantification. Cells undergoing DNA damage marked by γH2AX (red) in *gcn5[C137Y/C137Y]* (Q-Q’), *gcn5[E333st/E333st]* (R-R’) and *gcn5[E333st/C137Y]* (S-S’) as compared to wildtype (P-P’). Graphical representation of γH2AX positive cells undergoing DNA damage in the gcn5 mutants compared to the wildtype (T), n=24 for wildtype, n=54 for *gcn5[C137YC137Y]*, n=41 for *gcn5[E333st/E333st]*and n=31 for *gcn5[E333st/C137Y]* for quantification. Nuclei are stained with DAPI (Blue). n represents the number of individual primary lobes of the LG. Individual data points in the graphs represent individual primary lobes of the LG. Values are mean ± SD, and asterisk marks statistically significant differences (*p<0.05; **p<0.01; ***p<0.001, ****p<0.0001, Student’s t-test with Welch’s correction). Scale Bar: 50µm (A-S’). Nuclei are stained with DAPI (blue). Scale Bar: 50µm (A-S’).

### LG cellular subset-specific modulation of Gcn5 levels results in aberrant hematopoiesis

The observations from *gcn5* mutants led us to question if the modulation of Gcn5 levels in different cellular populations of the LG affects hematopoiesis. In order to modulate Gcn5 levels, we have depleted or over-expressed Gcn5 in different cellular subsets of the LG. Validation of *Hml-Gal4* mediated knockdown or overexpression of Gcn5 has been done using immunostaining with Gcn5 antibody (Fig S2A-C’). The FLAG-tagged Gcn5 overexpression construct has been validated using the anti-FLAG tag antibody by immunostaining (Fig S2D-E’).

Since the balance between the prohemocyte population and the differentiated blood cells is a determinant of the homeostasis conditions in the LG, we first depleted or overexpressed Gcn5 in the prohemocytes using *Dome-Gal4*. We observed that the PSC population remained unperturbed upon Gcn5 modulation in the prohemocytes as compared to wildtype (Fig 3A-C’ and M). However, the Dome positive progenitor pool decreased significantly upon Gcn5 knockdown whereas overexpression resulted in no significant change (Fig 3N). Over-expression of Gcn5 using *Dome-Gal4* led to a significant increase in both plasmatocyte and crystal cell differentiation whereas knockdown of Gcn5 did not affect blood cell differentiation compared to wildtype (Fig 3D-I’, O and P). An increase in DNA damage marked by Gamma H2Ax was observed upon knockdown of Gcn5 in the Dome population, whereas overexpression of Gcn5 did not lead to any change in DNA damage, as compared to wildtype (Fig 3J-L’ and Q). This data shows that elevated levels of Gcn5 in the prohemocytes promotes differentiation whereas knockdown has no effect on differentiation which could be due to Gcn5 mediated differential regulation of signaling in progenitors. Since PSC is known to produce signals that are required not only for maintaining the prohemocytes but also for priming them towards differentiation we modulated Gcn5 levels in the PSC using *Collier-Gal4*. Over-expression of Gcn5 in the PSC led to a cell-autonomous increase in PSC cell numbers whereas knockdown had no change in the PSC population as compared to the wildtype (Fig S3A-C’ and M). Plasmatocyte differentiation was reduced upon PSC-specific knockdown ofGcn5 whereas over-expression of Gcn5 did not affect plasmatocyte differentiation as compared to the wildtype (Fig S3D-F’ and N). Crystal cell differentiation remained unaffected upon modulation of Gcn5 in the PSC as compared to wildtype (Fig S3G-I’ and O). An increase in DNA damage was observed upon knockdown of Gcn5 in the PSC whereas over-expression did not lead to any change in DNA damage as compared to the wildtype (Fig S3J-L’ and P). These results indicate that Gcn5 modulation in the PSC has non-autonomous effects in the LG which could be due to abrogation of signals that are produced by the PSC which needs further investigation. Gcn5 over-expression in PSC results in an increase in PSC numbers which could be due to modulation of local signals that are required for regulating niche cell numbers which needs to be tested. The non-autonomous increase in DNA damage in the LG upon Gcn5 depletion in the PSC could be due to production of inflammatory signals by PSC cells upon Gcn5 depletion resulting in DNA damage which needs further investigation.

**Figure 3.**
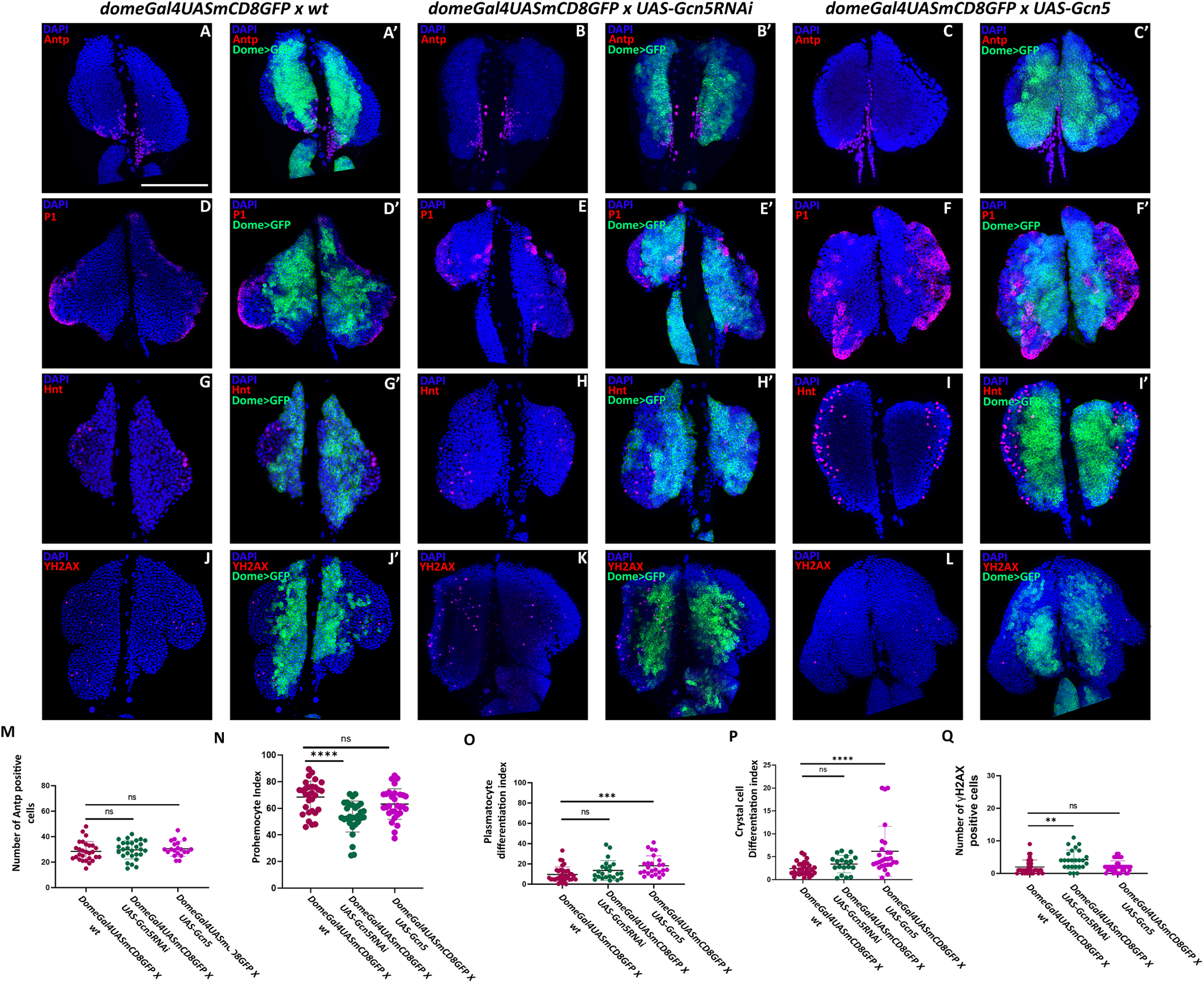
Modulation of Gcn5 levels in the prohemocytes alters blood cell homeostasis in the lymph gland. Posterior Signalling Center (PSC) population marked by Antennapedia (red) upon Dome-Gal4 mediated knockdown (B-B’) or over-expression (C-C’) of Gcn5 as compared to wildtype (A-A’). Plasmatocyte differentiation marked by P1 (red) upon Dome-Gal4 mediated knockdown (E-E’) or over-expression (F-F’) of Gcn5 as compared to wildtype (D-D’). Crystal cell differentiation marked by Hnt (red) upon Dome-Gal4 mediated knockdown (H-H’) or over-expression (I-I’) of Gcn5 as compared to wild type (G-G’). Cells undergoing DNA damage marked by γH2AX (red) upon Dome-Gal4 mediated knockdown (K-K’) or over-expression (L-L’) of Gcn5 as compared to wild type (J-J’). Graphical representation of PSC numbers upon modulation of Gcn5 levels in the Dome positive population compared to wildtype (M), n=28 for wildtype, n=28 for Gcn5 knockdown, and n=20 for Gcn5 over-expression. Graphical representation of prohemocyte index upon modulation of Gcn5 levels in dome population compared to wildtype (N), n=30 for wildtype, n=30 for Gcn5 knockdown, and n=30 for Gcn5 over-expression. Graphical representation of plasmatocyte differentiation index upon modulation of Gcn5 levels in dome population compared to wildtype (O), n=35 for wildtype, n=23 for Gcn5 knockdown, and n=25 for Gcn5 over-expression. Graphical representation of Crystal cell differentiation index upon modulation of Gcn5 levels in the dome population compared to wildtype (P), n=36 for wildtype, n=26 for Gcn5 knockdown, and n=34 for Gcn5 over-expression. Graphical representation of γH2AX positive cells upon modulation of Gcn5 levels in the dome population compared to wildtype (Q), n=33 for wildtype, n=27 for Gcn5 knockdown, and n=38 for Gcn5 over-expression. Nuclei are stained with DAPI (blue). Dome>GFP positive population (Green) indicates the expression domain of Dome-Gal4 activity in the LG. n represent the number of individual primary lobes of the LG. Individual data points in the graphs represent individual primary lobes of the Lymph Gland. Values are mean ± SD, and asterisk marks statistically significant differences (**p<0.01; ***p<0.001; ****p<0.0001, Student’s t-test with Welch’s correction). Scale Bar: 50µm (A-L’).

Since over-expression of Gcn5 in the prohemocyte population resulted in an increase in blood cell differentiation we wanted to understand if perturbing Gcn5 levels in the differentiated blood cell population using Hmldelta-Gal4 alters blood cell differentiation cell-autonomously. A non-cell-autonomous increase in the PSC population was observed upon knockdown or over-expression of Gcn5 using Hmldelta-Gal4 as compared to wildtype (Fig S4A-C’ and M). Plasmatocyte differentiation remained unaffected upon modulation of Gcn5 levels in the Hmldelta positive population compared to wildtype (Fig S4D-F’ and N), whereas a cell-autonomous increase in the crystal cell differentiation was observed upon overexpression of Gcn5 in the Hmldelta positive population whereas knockdown of Gcn5 did not affect the crystal cell differentiation compared to wildtype (Fig S4G-I’ and O). This data clearly indicates that Gcn5 has a cell autonomous role in regulating crystal cell differentiation. Now, Gcn5 is expressed in the crystal cells (Fig S5A-B) and cell autonomous overexpression of Gcn5 using Lozenge-Gal4, a crystal cell specific driver resulted in a significant increase in cell autonomous crystal cell differentiation demonstrating a cell-autonomous role (Fig S5C-D and G). An increase in DNA damage was observed upon knockdown or over-expression of Gcn5 levels in the Hmldelta population, compared to wildtype (Fig S4J-L’ and P). Overall, our analysis shows that modulation of Gcn5 levels in various sub-populations of the LG affects blood cell homeostasis.

### Prohemocyte-specific expression of Gcn5 domain deletion constructs alters the hematopoietic program and results in increased DNA damage

Gcn5 is an important component in the SAGA complex containing four different domains – the HAT, Pcaf, Ada, and Bromodomain (Fig.4A). We were further interested in investigating the functional role of each of the Gcn5 domains. In order to understand if any of the domain deletion constructs of Gcn5 perform a dominant negative function during hematopoiesis, we expressed Gcn5 lacking either the HAT, Pcaf, Ada or Bromodomain in the *Drosophila* prohemocytes using *tep4-Gal4*. A significant cell non-autonomous reduction in the PSC cell numbers was observed upon expression of Gcn5 domain deletion constructs in the prohemocyte population (Fig 4B-G). The tep4-GFP positive prohemocyte population was reduced upon expression of Bromo or Ada deletion constructs of Gcn5 using *tep4-Gal4*, whereas no change was observed upon expression of HAT or Pcaf domain deletion, compared to wildtype (Fig 4B’-F’ and H). A significant decrease in plasmatocyte differentiation was seen upon expression of Pcaf domain deletion construct of Gcn5, whereas, expression of Bromo or Ada domain deletion constructs of Gcn5 led to a significant increase in plasmatocyte differentiation. On the other hand, expression of Gcn5 lacking HAT domain using *tep4-Gal4* did not have any significant change in plasmatocyte differentiation (Fig 4B”-F” and I). An increase in total number of crystal cells was observed upon expression of all Gcn5 domain deletion constructs except Pcaf using tep4-Gal4 (Fig. 4B’’’-F’’’, 4J) whereas quantitation of the crystal cell differentiation index relative to the LG size showed that only Bromo domain deletion of Gcn5 had a statistically significant increase whereas expression of Gcn5 lacking HAT, Pcaf, or Ada domain using *tep4-Gal4* did not affect crystal cell differentiation, compared to wildtype (Fig 4B”’-F”’ and J). A significant increase in DNA damage was also observed upon expression of all domain deletion constructs of Gcn5 using *tep4-Gal4*, compared to wildtype (Fig S6A-F). These results indicate that various domains of Gcn5 play important roles during hematopoiesis which could be due to unique interactions facilitated by these domains. Expression of these domain deletion mutants in the wild type genetic background causes LG hematopoietic defects, suggesting that these domain deleted constructs may exert a dominant negative effect by competing with endogenous Gcn5 levels.

**Figure 4.**
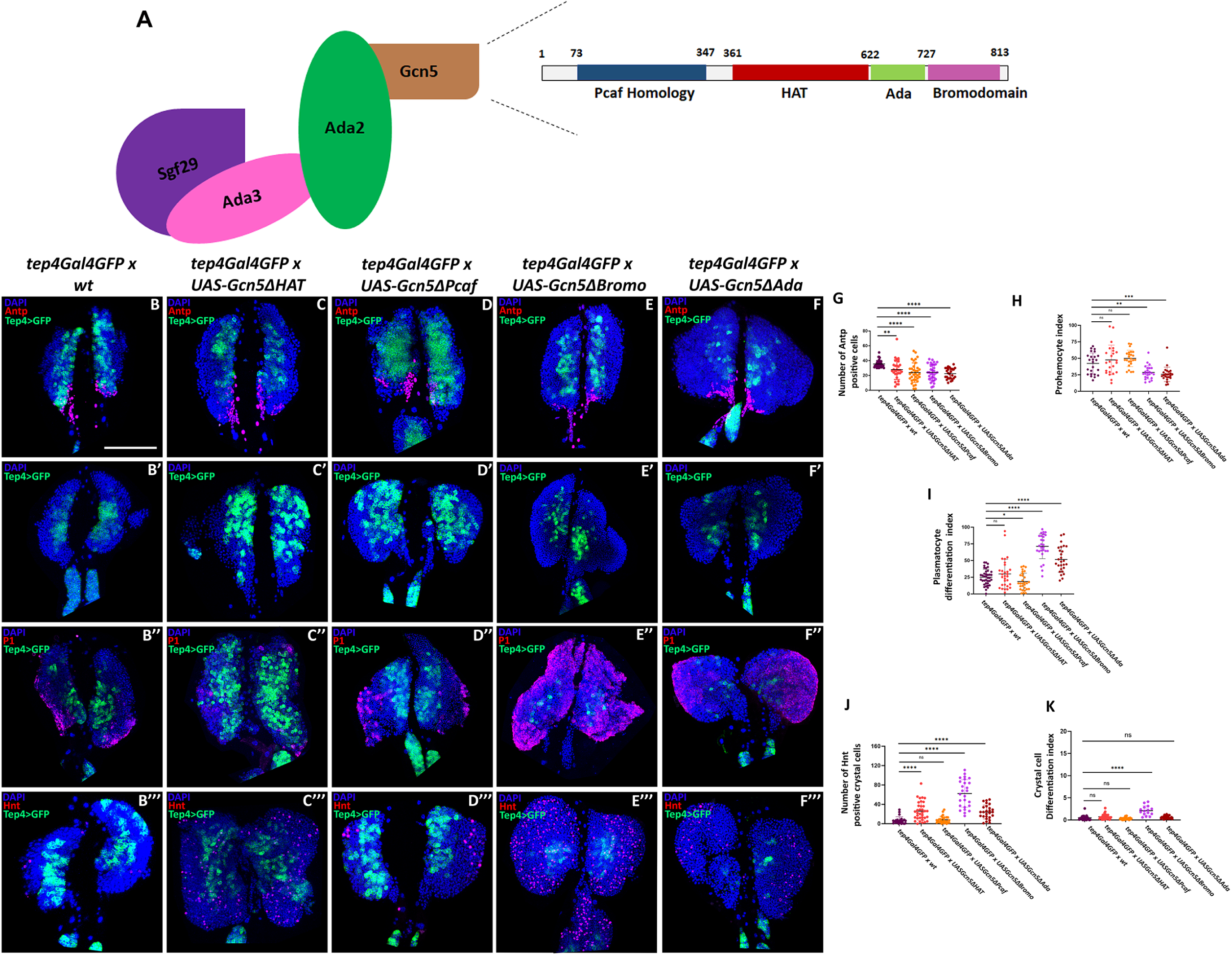
Prohemocyte-specific expression of Gcn5 domain deletion constructs affects LG homeostasis. Schematic of the SAGA HAT complex containing different modules – Sgf29, Ada2, Gcn5, and Ada3. Gcn5 is an 813 aa protein in Drosophila containing the Pcaf homology domain, HAT domain, Ada domain, and Bromodomain (A). Posterior Signalling Center (PSC) numbers marked by Antennapedia (red) upon expression of different Gcn5 domain deletion constructs in the tep-GFP (green) population using tep4-Gal4 (C-F), compared to wildtype (B). Progenitor index marked by tep (green) upon expression of different Gcn5 domain deletion constructs (C’-F’), compared to wildtype (B’). Plasmatocyte differentiation marked by P1 (red) upon expression of different Gcn5 domain deletion constructs in the tep-GFP population using tep4-Gal4 (green) (C”-F”), compared to wildtype (B”). Crystal cell differentiation marked by Hnt (red) upon expression of different Gcn5 domain deleted constructs in the tep-GFP population using tep4-Gal4 (green) (C”’-F”’), compared to wildtype (B”’). Graphical representation of PSC numbers upon expression of Gcn5 domain deletion constructs, compared to wildtype (G), n=40 for wildtype, n=29 for Gcn5ΔHAT expression, n=35 for Gcn5ΔPcaf expression, n=33 for Gcn5ΔBromo expression, and n=26 for Gcn5ΔAda expression in the tep population. Graphical representation of Prohemocyte index upon expression of Gcn5 domain deletion constructs, compared to wildtype (H), n=24 for wildtype, n=24 for Gcn5ΔHAT expression, n=34 for Gcn5ΔPcaf expression, n=24 for Gcn5ΔBromo expression, and n=24 for Gcn5ΔAda expression in the tep population. Graphical representation of Plasmatocyte differentiation index upon expression of Gcn5 domain deleted constructs, compared to wildtype (I), n=38 for wildtype, n=30 for Gcn5ΔHAT expression, n=29 for Gcn5ΔPcaf expression, n=27 for Gcn5ΔBromo expression, and n=27 for Gcn5ΔAda expression in the tep population. Graphical representation of Crystal cell numbers upon expression of Gcn5 domain deleted constructs, compared to wildtype (J), n=23 for wildtype, n=35 for Gcn5ΔHAT expression, n=26 for Gcn5ΔPcaf expression, n=26 for Gcn5ΔBromo expression, and n=26 for Gcn5ΔAda expression in the tep population. Graphical representation of Crystal cell Differentiation index upon expression of Gcn5 domain deleted constructs, compared to wildtype (K), n=20 for wildtype, n=34 for Gcn5ΔHAT expression, n=25 for Gcn5ΔPcaf expression, n=15 for Gcn5ΔBromo expression, and n=27 for Gcn5ΔAda expression in the tep population. Nuclei are stained with DAPI (blue). n represent the number of individual primary lobes of the LG. Individual data points in the graphs represent individual primary lobes of the LG. Values are mean ± SD, and asterisk marks statistically significant differences (*p<0.05; **p<0.01; ***p<0.001; ****p<0.0001, Student’s t-test with Welch’s correction). Scale Bar: 50µm (B-F’’’).

### Autophagic flux in the *Drosophila* blood cells is negatively regulated by Gcn5

Increasing evidence suggests that Gcn5 can regulate gene expression programs linked to cellular metabolism ^65^. Autophagy is a cellular process that is closely associated with cell metabolism regulation and intracellular quality control as it is involved in the turnover of undesired cellular components, damaged or dysfunctional organelles^24,25,66^. Autophagic flux is regulated by a number of molecules, one of them is Transcription Factor EB (TFEB) which is a known non-histone acetylation target of Gcn5. TFEB regulates expression of genes related to autophagosome formation and lysosome production which are critical for autophagy. Mammalian cell culture-based studies show that acetylation of TFEB by Gcn5 decreases TFEB transcriptional activity by disrupting its dimerization thereby affecting autophagy. Studies done in the *Drosophila* fat body indicate that Gcn5 negatively regulates formation of Atg8a puncta in starved conditions ^59^.

Hence, we sought to determine if Gcn5 regulates autophagic flux in *Drosophila* blood cells. We genetically modulated Gcn5 levels using a pan hemocyte driver, *Hml-Gal4* and probed the LG hemocytes for Ref(2)P/p62, an adaptor protein that is itself cleared during autophagy and Atg8 (homologue of human LC3). Our results indicate that there are no p62 puncta upon knockdown of Gcn5, whereas over-expression of Gcn5 led to an increase in p62 puncta, compared to wildtype (Fig 5A-C, D). On the contrary, immunostaining with Atg8 showed an increase in Atg8 puncta upon knockdown of Gcn5, whereas over-expression of Gcn5 leads to a significant reduction in the Atg8 puncta compared to wildtype (Fig 5A’-C’, E). We also looked at the transcript levels of a panel of Atg genes involved during autophagy in the hemocytes upon over-expression of Gcn5 and our results indicate that Gcn5 over-expression using pan hemocyte driver, *Hml-Gal4* leads to a reduction in transcript levels of a panel of Atg genes in the hemocytes (Fig S7).

**Figure 5.**
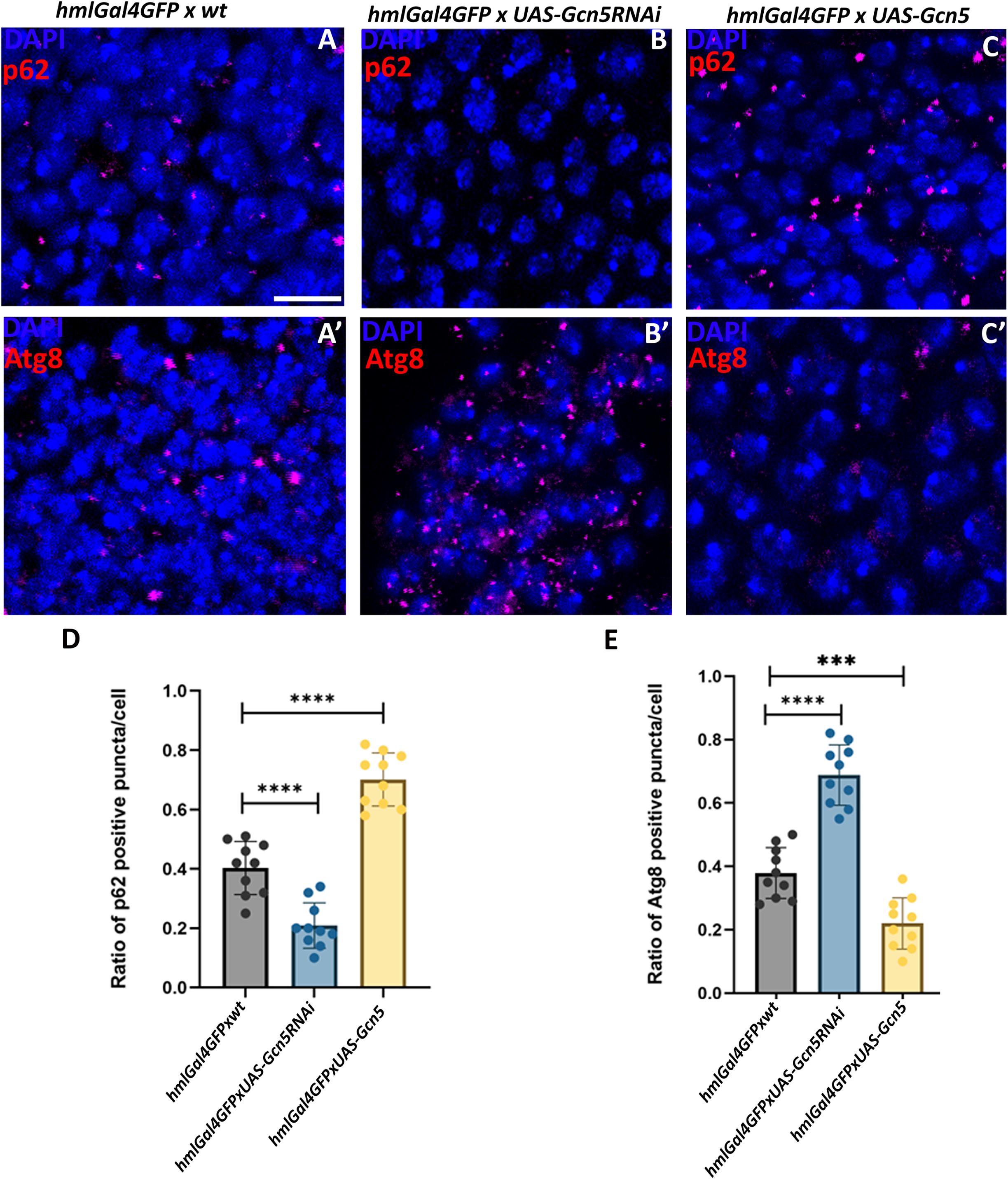
Autophagic flux in *Drosophila* blood cells is negatively regulated by Gcn5. p62 (red) labeling the autophagy adaptor protein upon Hml-Gal4 mediated Gcn5 knockdown (B), or overexpression (C) as compared to wildtype (A). Atg8 (red), the autophagosomal marker upon Hml-Gal4 mediated Gcn5 knockdown (B’) or over-expression (C’) as compared to wildtype (A’). Nuclei were stained with DAPI (blue). Graphical representation of the p62 positive puncta/cell (D). Graphical representation of Atg8 positive puncta/cell (E). Values are mean ± SD, and asterisk marks statistically significant differences (***p<0.001; ****p<0.0001, Student’s t-test with Welch’s correction). Scale Bar: 20µm (A-E’).

### Genetic and chemical ablation of autophagy boosts blood cell differentiation in the primary LG lobe

Autophagy has been shown to regulate cellular homeostasis of all hematopoietic lineages in mammals^67^. Mis-regulation of autophagy in hematopoietic stem cells (HSC) leads to dysfunction and loss of HSCs with age^27^. Transcription factors that are considered to be master regulators of hematopoiesis have been shown to transcriptionally control autophagy related genes^29,68^. In *Drosophila*, Atg6 has been shown to play a role in multiple vesicle trafficking pathways and hematopoiesis^32^.

Since we found that Gcn5 negatively regulates autophagy in *Drosophila* hemocytes, we asked the question whether key autophagy related genes regulate LG hematopoiesis. Since Gcn5 regulates autophagy via TFEB, a key regulator in autophagosome and lysosome biogenesis, we perturbed TFEB and several autophagy effector genes like Atg8a, Atg5, or Atg18a specifically in the *Drosophila* prohemocytes using tep4-Gal4. We observed that depleting TFEB or the autophagy effector genes in the tep population led to a drastic increase in both plasmatocyte and crystal cell differentiation in the LG along with a decrease in the tep4 positive prohemocyte index (Fig 6A-L, Fig S8M). Atg8 or TFEB knockdown did not affect the niche cell numbers, however Atg5 resulted in a significant increase in niche numbers whereas Atg18a knockdown resulted in a significant decrease. Depleting TFEB or the autophagy effector genes in the tep population led to a drastic increase in DNA damage. (Fig S8A-L). In addition to the genetic perturbation-based analysis, chemical inhibition of autophagy by treating the larvae with autophagy inhibitor, chloroquine (CQ) for 16 hours resulted in a significant increase in blood cell differentiation in the LG (Fig 6M-R). We also validated the effect of CQ on autophagy and we find that there is an increase in p62 positive puncta and a decrease in Atg8 positive puncta per cell in the *Drosophila* LG upon CQ treatment for 16 hours (Fig S9A-F).

**Figure 6.**
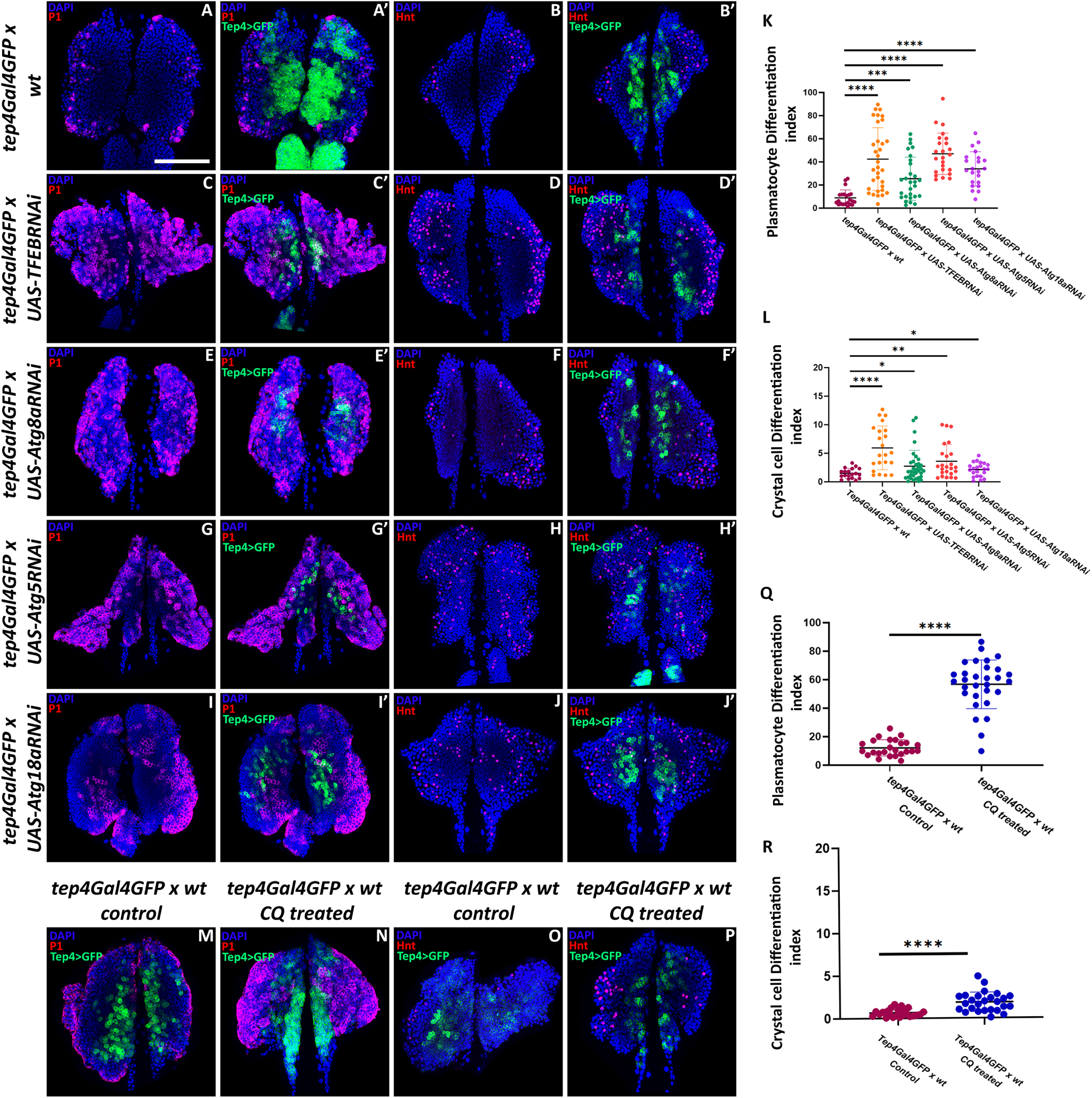
Genetic and chemical ablation of autophagy leads to aberrant blood cell differentiation. Plasmatocyte differentiation marked by P1 (red) upon tep4-Gal4 mediated knockdown of TFEB (C-C’), Atg8a (E-E’), Atg5 (G-G’), or Atg18 (I-I’) in the tep population (green) as compared to wildtype (A-A’). Crystal cell differentiation marked by Hnt (red) upon tep4-Gal4 mediated knockdown of TFEB (D-D’), Atg8a (F-F’), Atg5 (H-H’), or Atg18 (J-J’) in the tep population (green) as compared to wildtype (B-B’). Graphical representation of Plasmatocyte differentiation index upon knockdown of TFEB (n=33), Atg8a (n=28), Atg5 (n=24), or Atg18 (n=22) in the tep population, compared to the wildtype (n=22) (K). Graphical representation of Crystal cell differentiation index upon knockdown of TFEB (n=22), Atg8a (n=35), Atg5 (n=24), or Atg18 (n=22) in the tep population, compared to the wildtype (n=19) (L). Plasmatocyte differentiation(red) upon Chloroquine treatment (N) as compared to control (M). Crystal cell differentiation upon Chloroquine treatment (P) as compared to control (O). Graphical representation of Plasmatocyte Differentiation Index upon Chloroquine treatment (n=30), compared to control (n=25) (Q). Graphical representation of Crystal Cell differentiation index upon Chloroquine treatment (n=26), compared to control (n=23) (R). Nuclei are stained with DAPI (blue). n represents the number of individual primary lobes of the LG. Individual data points in the graphs represent individual primary lobes of the LG. Values are mean ± SD, and asterisk marks statistically significant differences (*p<0.05; ***p<0.001; ****p<0.0001, Student’s t-test with Welch’s correction). Scale Bar: 50µm (A-P).

### Chemical or genetic modulation of mTORC1 activity controls blood cell differentiation

mTOR signaling, specifically mTORC1 has been shown to play a significant role in hematopoietic lineage commitment in mice^69^. mTORC1 acts as a master regulator of autophagy and controls various steps of the autophagic process^26^. In nutrient replete conditions, mTORC1 phosphorylates ULK1, a key initiator kinase in autophagy thereby disrupting its interaction with AMPK and ULK1 activation by AMPK^70–73^. Similarly, mTORC1 phosphorylates TFEB in nutrient availability conditions at residues -S142 and S211 subjecting TFEB to localize in the cytoplasm rendering it inactive^74^. Since mTORC1 regulates critical nodal points that control autophagy we were curious to know if modulating mTORC1 activity alters blood cell homeostasis in the LG. We modulated mTORC1 activity by chemical modulation. We used 3BDO for mTOR activation and Rapamycin for mTOR inhibition. Upon treatment with the mTOR activator – 3BDO for 16 hours, a significant increase in both plasmatocyte and crystal cell differentiation was observed as compared to the control treatment (Fig 7A-F). Similarly, activation of mTOR genetically by overexpressing Rheb using tep4-Gal4 resulted in an increase in both plasmatocyte and crystal cell lineage differentiation (Fig S10D-H and J). Whereas treatment with mTOR inhibitor – Rapamycin for 16 hours led to a decrease in plasmatocyte and crystal cell differentiation, compared to the control treatment (Fig 7G-L). While genetically depleting Tor did not result in any significant change in plasmatocyte and crystal cell differentiation, Raptor depletion resulted in a decrease in crystal cell differentiation whereas no significant change was observed with respect to plasmatocyte differentiation (Fig S10A-B, E-F and I). Also, Tep4 progenitor specific knockdown of Tor and Raptor did not have any non-cell autonomous effect on niche numbers. On the other hand, mTOR activation via Rheb resulted in a non-cell autonomous decrease in niche numbers as compared to the wildtype (Fig S11A-D, I). Abrogation of mTOR signaling by tep4-Gal4 mediated depletion of Tor resulted in a decrease in γ-H2Ax positive puncta in the LG whereas Raptor depletion or Rheb over-expression did not have any effect as compared to the wild type control (Fig S11E-H, J).

**Figure 7.**
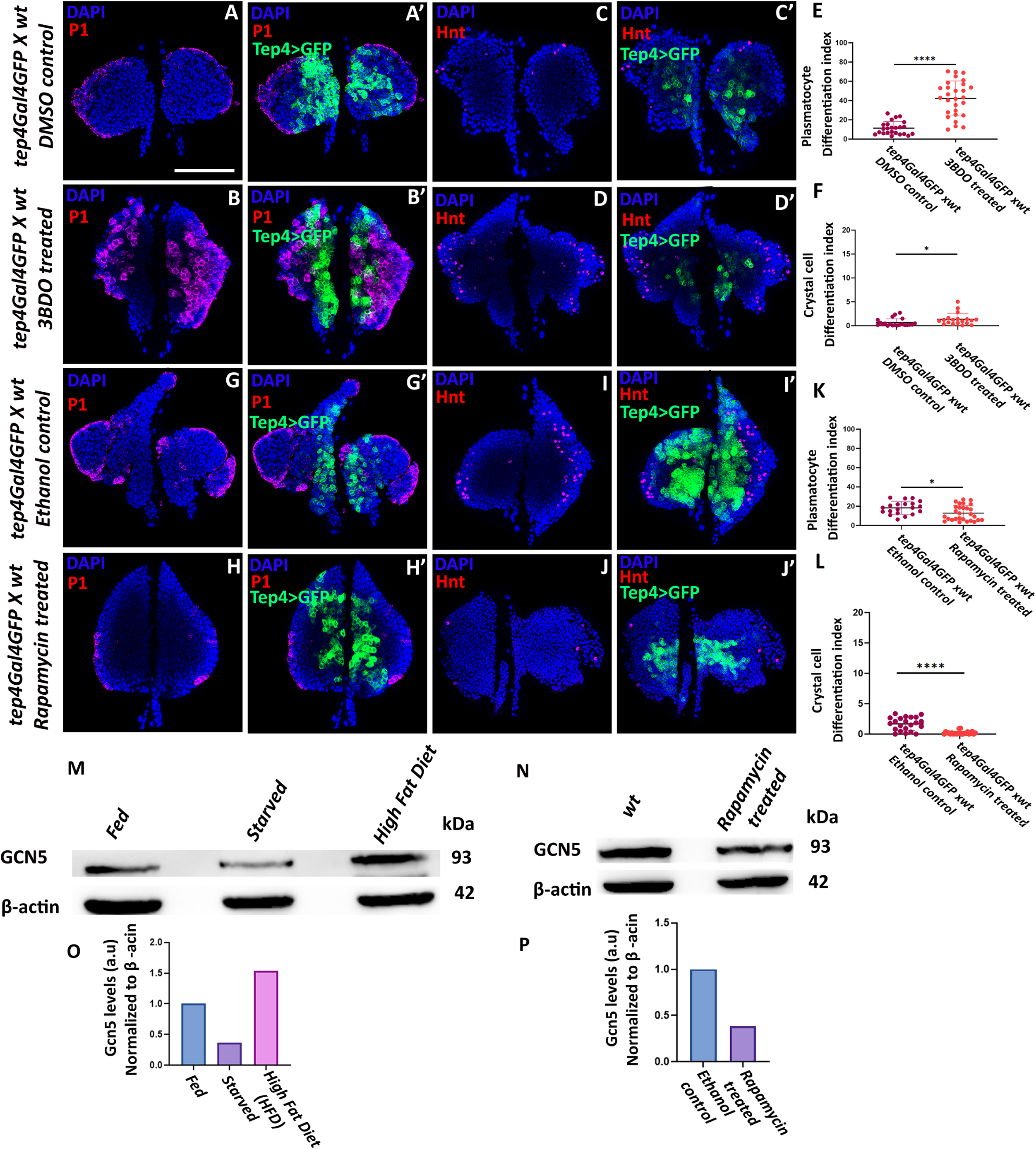
Modulation of mTORC1 activity regulates blood cell differentiation in the Lymph gland. Plasmatocyte differentiation marked by P1 (red) upon 3BDO treatment (B-B’) as compared to control (A-A’). Crystal cell differentiation marked by Hnt (red) upon 3BDO treatment (D-D’) compared to control (C-C’). Graphical representation of Plasmatocyte Differentiation Index upon 3BDO treatment (n=28) compared to control (n=24) (E). Graphical representation of Crystal cell differentiation index upon 3BDO treatment (n=19) compared to control (n=21) (F). Plasmatocyte differentiation marked by P1 (red) upon Rapamycin treatment (H-H’) compared to control (G-G’). Crystal cell differentiation marked by Hnt (red) upon Rapamycin treatment (J-J’) compared to control (I-I’). Graphical representation of Plasmatocyte Differentiation Index upon Rapamycin treatment (n=28) compared to control (n=20) (K). Graphical representation of Crystal cell differentiation index upon Rapamycin treatment (n=23) compared to control (n=22) (L). Immunoblot showing GCN5 (93 kDa) protein levels in different diet conditions – fed, starved, and high-fat diet (M). Immunoblot showing GCN5 (93 kDa) protein levels upon Rapamycin treatment compared to wildtype (N). β-actin (42kDa) was used as the loading control. Nuclei are stained with DAPI (blue). Quantification of GCN5 protein levels in the fed, starved and High Fat diet scenario (O). Quantification of GCN5 protein levels in the wildtype as compared to rapamycin treatment (P). n represent the number of individual primary lobes of the LG. Individual data points in the graphs represent individual primary lobes of the LG. Values are mean ± SD, and asterisk marks statistically significant differences (*p<0.05; **p<0.01; ***p<0.001; ****p<0.0001, Student’s t-test with Welch’s correction). Scale Bar: 50µm (A-J’).

### Gcn5 acts as a nutrient sensor and is regulated by mTORC1

Enzymatic activity of Gcn5 is regulated by the cellular and metabolic energy state which in turn controls gene expression programs^65^. Gcn5 is capable of sensing Acetyl Co-A which is a central metabolic intermediate^75^. Since Gcn5 levels are critical in controlling gene expression programs and transcriptional noise^76^, we tested if Gcn5 levels are responsive to altered nutrient conditions. We subjected *Drosophila* larvae to different dietary conditions viz. Fed, starved and high fat diet (HFD) and probed for Gcn5 levels in these conditions. Our results indicate that there is a decrease in Gcn5 levels upon starvation whereas an increase in high fat diet (HFD) conditions (Fig 7M, O) Gcn5 levels were normalized to β-actin in the fed, starved and high diet scenario (Fig 7O) and upon Rapamycin treatment (Fig 7P). mTOR is known to control protein translation in response to nutrient stress signals; under nutrient deprivation, mTOR is inhibited thereby inducing autophagy. There are few studies that have linked mTOR to histone acetyltransferase Gcn5 and therefore nutrient response^59,77^ . To further determine if Gcn5 levels are regulated by mTORC1 activity, we treated larvae with mTOR inhibitor Rapamycin and probed for Gcn5 levels. We found that there was a decrease in Gcn5 levels upon mTOR inhibition as compared to control (Fig 7N, P).

### mTORC1 overrides the effect of Gcn5 in regulating blood cell homeostasis in the LG

mTORC1 regulates autophagy at multiple nodal points for example by phosphorylation of initiator kinase ULK1 or by phosphorylating TFEB preventing its nuclear translocation. On the other hand, Gcn5 regulates autophagy by acetylation of TFEB preventing its dimerization and nuclear translocation thereby inhibiting autophagy^59^. Since both mTORC1 and Gcn5 converge on and exert their effect over TFEB by phosphorylation or acetylation respectively, we wanted to investigate if one can override the effector function of the other. Since our results indicate that mTOR activity can regulate Gcn5 levels, we wanted to test if mTORC1 can override the effect of Gcn5 modulation in controlling LG hematopoiesis. Activation of mTOR by treatment with 3BDO in tep4-Gal4 mediated prohemocyte specific Gcn5 knockdown genetic background led to increased plasmatocyte and crystal cell differentiation as compared to the tep4-Gal4 mediated prohemocyte specific Gcn5 knockdown control (Fig 8A-D’, I and J). Similarly, inhibition of mTOR by Rapamycin treatment in the dome-Gal4 mediated prohemocyte specific Gcn5 over-expression genetic background resulted in a significant reduction in plasmatocyte and crystal cell differentiation as compared to the dome-Gal4 mediated prohemocyte specific Gcn5 over-expression control (Fig 8E-H’, K and L). This over-riding effect is based on chemical intervention data and would need further genetic experiments to fully establish the regulatory role of mTORC1. Taken together, our results demonstrate that a fine balance between mTORC1 mediated phosphorylation of TFEB and Gcn5 mediated acetylation of TFEB maintains the autophagic flux. Nutrient conditions, mTORC1 activity and Gcn5 levels are key determinants of autophagic flux in the LG. Maintenance of the autophagic flux and turnover is critical for regulating blood cell homeostasis in the LG. Thus, Gcn5-mTOR-TFEB signaling axis controls autophagy thereby regulating *Drosophila* blood cell homeostasis in the LG (Fig 9).

**Figure 8.**
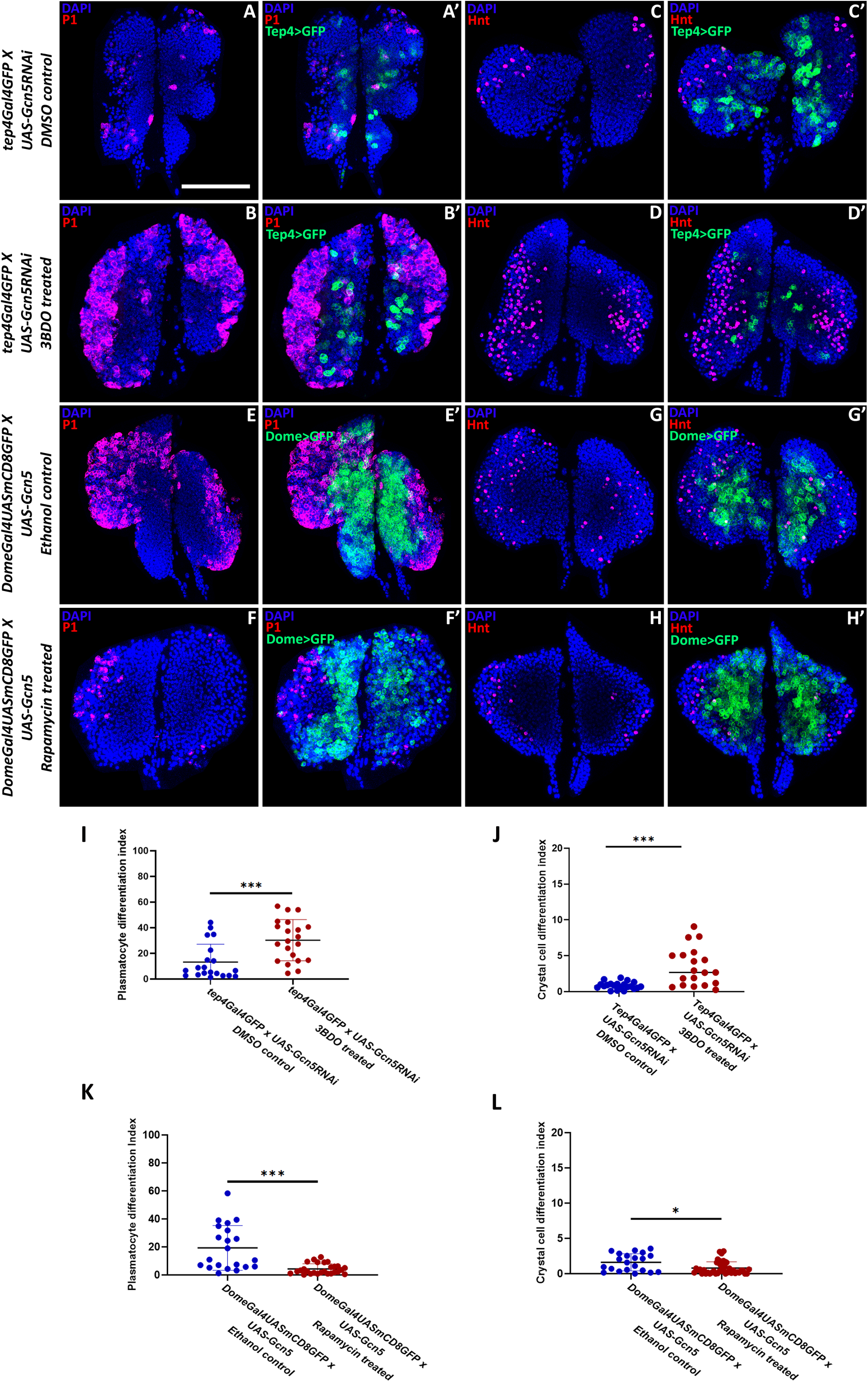
mTORC1 can override the effect of Gcn5 level modulation over autophagy. Plasmatocyte differentiation marked by P1 (red) upon 3BDO treatment in tep4-Gal4 mediated (green) Gcn5 knockdown background (B-B’) compared to control treatment in the same genetic background (A-A’). Crystal cell differentiation marked by Hnt (red) upon 3BDO treatment in the tep-mediated (green) Gcn5 knockdown background (D-D’) compared to control treatment in the same genetic background (C-C’). Plasmatocyte differentiation marked by P1 (red) upon Rapamycin treatment in Dome-mediated (green) Gcn5 over-expression background (F-F’) compared to control treatment in the same genetic background (E-E’). Crystal cell differentiation marked by Hnt (red) upon Rapamycin treatment in Dome-Gal4 mediated (green) Gcn5 over-expression background (H-H’) as compared to control treatment in the same genetic background (G-G’). Graphical representation of Plasmatocyte Differentiation Index upon 3BDO treatment (n=21) and control treatment (n=20) in the tep-mediated Gcn5 knockdown background (I). Graphical representation of Crystal cell differentiation index upon 3BDO treatment (n=19) and control treatment (n=21) in the tep-mediated Gcn5 knockdown background (J). Graphical representation of Plasmatocyte Differentiation Index upon Rapamycin treatment (n=27) and control treatment (n=21) in the Dome-mediated Gcn5 over-expression background (K). Graphical representation of Crystal Cell differentiation index upon Rapamycin treatment (n=43) and control treatment (n=21) in the Dome-mediated Gcn5 over-expression background (L). Nuclei are stained with DAPI (blue). n represent the number of individual primary lobes of the LG. Individual data points in the graphs represent individual primary lobes of the LG. Values are mean ± SD, and asterisk marks statistically significant differences (**p<0.01; ***p<0.001, Student’s t-test with Welch’s correction). Scale Bar: 50µm (A-H’).

**Figure 9:**
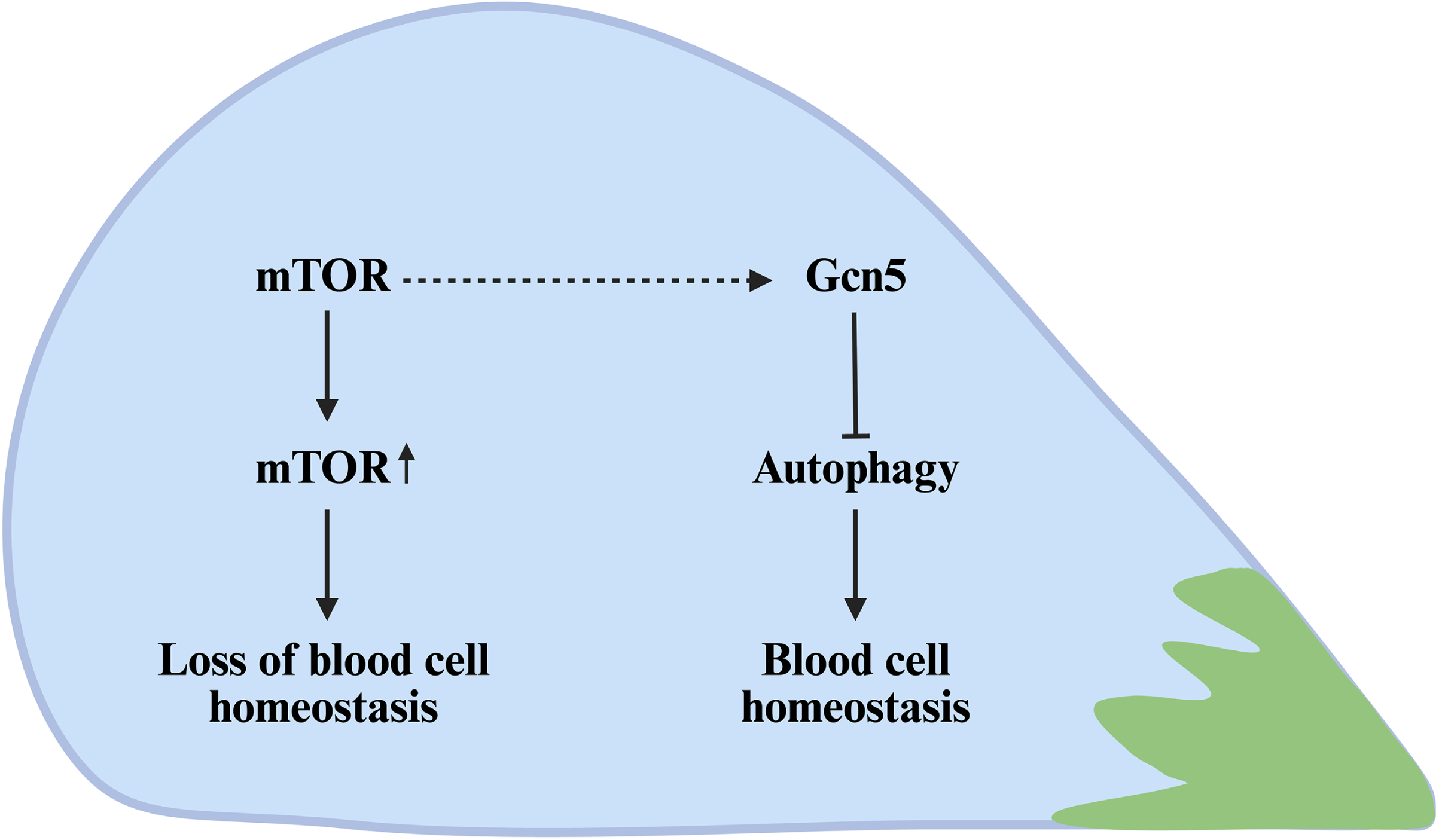
Gcn5-mTORC1 mediated autophagy regulation controls blood cell homeostasis. A cartoon summarizing how Gcn5-mTORC1-signaling axis regulates autophagy to control blood cell homeostasis in the Drosophila LG. Lab, R. (2026). Figure 9. Gcn5-mTORC1 mediated autophagy regulation controls blood cell homeostasis. Created in BioRender. https://BioRender.com/lq2uud3

## Discussion

Gcn5 is highly expressed in acute myeloid leukemia (AML) and helps in the maintenance of immature leukemic blasts ^54,55^. While its function in the context of leukemia is well studied, Gcn5 function during physiological hematopoiesis is unexplored. Here, we explore Gcn5 role during normal physiological hematopoiesis in *Drosophila*. Our work shows that Gcn5, a histone acetyltransferase plays an important role during *Drosophila* LG hematopoiesis. We find that Gcn5 is expressed in all the LG cellular subsets and its levels are critical for controlling hematopoiesis. Structure-function analysis of Gcn5 shows that expression of Gcn5 lacking any of its constituent domains in wild type genetic background results in a dominant negative phenotype in the LG. Our results indicate that optimum levels of Gcn5 are critical in the prohemocytes as excess Gcn5 results in increased blood cell differentiation. We find that the number of PSC cells are altered either in an autonomous or non-autonomous manner where PSC-specific or non-PSC-specific modulation of Gcn5 levels results in changes in niche cell numbers which could be due to perturbation of signaling pathways like Insulin, mTOR, Dpp which are critical in maintaining the niche cells ^21,78^ which needs further exploration. Our results also suggest that niche cells numbers are reduced in the *gcn5* null mutants and transheterozygotes of *gcn5* mutant alleles which we speculate could be due to abrogation of local niche supporting pathways such as mTOR signaling, which is known to be a critical regulator of PSC maintenance. Since these are whole animal mutants there could be a number of signals that could be perturbed which needs to be investigated. Our results show that there is increased differentiation in *gcn5* mutant alleles and transheterozygotes which could be due to abrogation of various signaling pathways that might be affected systemically and LGs are highly sensitive to systemic signals which could result in aberrant hematopoiesis which needs further investigation. Also, we observe that PSC numbers are modulated upon perturbation of Gcn5 using *HmlΔ-Gal4,* uncovering a non-cell autonomous effect on the niche which would need further investigation to determine if it represents a feedback mechanism. Further, we find that Gcn5 acts as a nutrient sensor and its levels are critical for controlling the autophagic flux in the LG. In order to investigate whether autophagy players regulate LG hematopoiesis, we carried out a targeted genetic analysis of key autophagy regulators and our results indicate that disrupting autophagy abrogates LG hematopoiesis. Increased Gcn5 levels in the hemocytes led to a decrease in the autophagic flux whereas depleting Gcn5 resulted in increased autophagy. LG phenotypes upon genetic modulation of Gcn5 corroborate with the phenotypes obtained upon perturbation of autophagy regulators, providing a potential parallel mechanism.

Since mTORC1 regulates autophagy in response to nutrient conditions, we investigated whether modulating mTORC1 activity affects hematopoiesis. Our data elucidates that modulating mTORC1 activity regulates hematopoiesis and these effects are consistent with observations obtained upon genetic modulation of autophagy. Inhibiting mTORC1 activity using Rapamycin results in Gcn5 down-regulation, suggesting a potential regulatory connection between mTORC1 signaling and Gcn5 expression. One of the common molecular link through which both mTORC1 and Gcn5 could potentially regulate autophagy is through Transcription factor EB (TFEB), a member of the microphthalmia-associated transcription factor/transcription factor E (MiTF/TFE) family^79^ . Our results demonstrate that mTORC1 can override Gcn5 in regulating LG hematopoiesis. Although we show this over-ride effect with chemical intervention, further genetic epistasis-based analysis will further strengthen this data providing important insights. Overall, our data points to a model where Gcn5 and mTORC1 control autophagy to regulate *Drosophila* LG hematopoiesis.

Expression analysis of Gcn5 in the LG indicates that Gcn5 is expressed in all the LG cellular sub-populations. *gcn5* mutant alleles display loss of blood cell homeostasis. Increase in plasmatocyte or crystal cell differentiation could be due to a contribution of systemic signals that are altered upon loss of Gcn5 in the entire organism as the LG is capable of responding to external stimuli^80^ . *gcn5* mutant alleles either in homozygous or trans-heterozygous conditions show an increased accumulation of DNA damage indicative of its role in DNA damage repair pathways^49,81^. Prohemocyte-specific over-expression of Gcn5 leads to increased plasmatocyte as well as crystal cell differentiation whereas depletion of Gcn5 levels has no significant effect on differentiation. *Hmldelta-Gal4* mediated Cortical Zone (CZ) specific over-expression of Gcn5 resulted in a cell autonomous increase in crystal cell differentiation whereas Gcn5 knockdown had no effect. These results suggest that Gcn5 acts in a lineage-specific manner cell-autonomously in the differentiated hemocytes. Higher levels of Gcn5 skews the balance towards increased blood cell differentiation. Higher Gcn5 expression has been observed previously in many carcinomas and leukemia promoting cell proliferation, growth and cancer progression ^50–55^. Structure-function analysis shows that expression of Gcn5 lacking the Bromo or Ada domain in the wild type genetic background results in a dominant negative phenotype in terms of the maintenance of prohemocyte population itself and plasmatocyte differentiation whereas expression of Gcn5 lacking HAT or Bromo or Ada results in a dominant negative effect on crystal cell differentiation indicating that the constituent domains of Gcn5 might be playing specific functions in regulating various aspects of LG hematopoiesis via specific interacting partners. The constituent domains in Gcn5 could be performing unique functions in regulating homeostasis based on their interacting partners which needs to be explored. Gcn5 responds to changes in cellular energetic and metabolic states ^65^. Gcn5 via its non-histone acetylation target, Transcription factor EB (TFEB) is capable of regulating autophagy which is directly linked to the nutritional status of the cell. Since Gcn5 is capable of sensing nutrient signals, its role in regulating the sensing by blood progenitors needs to be studied in detail and if it has any other potential crosstalk with signaling pathways that act downstream of nutrient and metabolic signals. TFEB acetylation by Gcn5 is inhibitory for TFEB as it prevents its dimerization and hence nuclear translocation. Upon nuclear translocation, TFEB activates genes responsible for autophagosome and lysosome biogenesis^59^ .

In order to probe whether Gcn5 mediated regulation is due to its control over autophagy, we probed whether perturbing Gcn5 has any effect on the autophagic flux in the hemocytes. Our results suggest that depletion of Gcn5 leads to an upregulation of autophagy increasing the flux whereas over-expression of Gcn5 results in downregulation of autophagy lowering the autophagic flux. These observations were intriguing as this establishes an important role of autophagy in maintaining LG homeostasis. We investigated this further to find out if there is a direct link between autophagy regulators (Atg genes) including TFEB in controlling LG hematopoiesis. Genetic perturbation of the autophagy pathway regulators or TFEB in the prohemocytes consistently accelerated blood cell differentiation. Chemical inhibition of autophagy induced by Chloroquine also recapitulates this phenotype. There is literature indicating a critical role for autophagy in development of the blood system, self-renewal of HSCs and their mobilization^27,30,31,35^

Our observations on the role of autophagy in *Drosophila* LG hematopoiesis support these findings. We then moved on to test if mTORC1 could regulate hematopoiesis. Chemical or genetic modulation of mTORC1 activity which is a master molecule that regulates autophagy alters LG hematopoiesis. mTORC1 activity regulates Gcn5 levels as inhibition of mTORC1 activity resulted in lower levels of Gcn5.

mTORC1 regulates autophagy via two arms -One via the initiator kinase, Ulk1 in response to either nutrient depletion or replete conditions via inhibitory phosphorylation of TFEB^71–73^ . TFEB phosphorylation by mTORC1 subjects it to degradation thereby inhibiting autophagy^82^. Since TFEB is the common link that is regulated by both mTORC1 and Gcn5, we wanted to understand whether one is epistatic to the other in controlling hematopoiesis. We find that mTORC1 can override Gcn5 in controlling hematopoiesis by chemical intervention approach. Although, further genetic epistasis experiments will be needed to show that this effect is indeed by acting upstream and controlling Gcn5. We speculate that during physiological hematopoiesis a balance between phosphorylation and acetylation of TFEB mediated by mTORC1 and Gcn5 respectively regulates levels of autophagy which needs further experimental investigation. Perturbation of Gcn5 or mTORC1 activity could disturb this balance thereby deregulating hematopoiesis. Our findings shed light on the importance of non-histone acetylation function of Gcn5 in hematopoiesis. This could especially be important as the LG prohemocytes are known to respond to cell intrinsic and extrinsic nutrient signals^20^. The nutrient state could regulate autophagy thereby dictating the differentiation trajectory of the precursor cells. Since Gcn5 is highly expressed in acute myeloid leukemia^54,55^ it becomes important to understand how Gcn5 level is regulated and what cellular processes Gcn5 is capable of regulating in order to control tissue homeostasis. Our work sheds novel insights on the role of Gcn5 during normal hematopoiesis. Our results demonstrate that organismal Gcn5 levels fluctuate in response to dietary challenges. Gcn5 could be acting as a downstream effector that translates systemic metabolic changes to modulate autophagic flux in blood cells. It would be interesting to understand the kind of nutrient signals to which Gcn5 is capable of responding and whether Gcn5 has any crosstalk with other nutrient sensing molecules and pathways other than mTORC1. The nutrient sensing function of Gcn5 given the ability of hemocytes to respond to nutrient cues warrants further investigation. We show that Gcn5 fine tunes autophagy in conjunction with mTORC1 where mTORC1 activity is capable of regulating Gcn5 and can override its function in modulating autophagy. Our findings shed an important and broad role of Gcn5 and mTORC1 in controlling autophagy that ultimately regulates *Drosophila* hematopoiesis (Figure 9).

## Materials and Methods

List of *Drosophila* stocks, antibodies used for immunofluorescence-based experiments have been listed in a detailed manner in the Supplemental Information (SI). The experimental protocol for lymph gland dissection, immunohistochemistry and image analysis has been mentioned in the SI. The procedure for protein extraction, western blotting and quantitative real time PCR has been mentioned in the SI along with the list of primers used for qRT experiments. The chemical treatment procedure has been described in a detailed manner in the SI. Statistical analysis performed for each of the experiments has been described in the statistical analysis section in the SI.

## Data, Materials, and Software availability

All the data related to this study are included in the article and/or in the SI.

## Acknowledgements

We thank the Bloomington Drosophila Stock Center, Developmental Studies Hybridoma Bank and the fly community for fly stocks and antibodies. We would particularly like to thank Clement Carre, Lucas Waltzer, Mohit Prasad and Bhupendra Shravage for help with fly stocks. We would like to thank the core central facilities and the technical support staff at ACTREC – Tata Memorial Centre. We also thank our lab members for critical discussions. This work was funded by Basic and Translational Research in Cancer grant (No.1/3(7)/2020/TMC/R&D-II/8823 Dt.30.07.2021), Capacity Building and Development of Novel and Cutting-edge Research Activities (No.1/3(4)/2021/TMC/R&D-II/15063 Dt.15.12.2021) from Department of Atomic Energy (DAE), Government of India.

## Author contributions

Conceptualization: AAR and RJK; Methodology: AAR, MK, SM and LK; Validation: AAR, MK; Formal analysis: AAR, MK, SM and LK; Investigation: AAR, MK, SM and LK; Resources: RJK; Data curation: AAR, MK; Writing - original draft: AAR, MK RJK; Writing - review & editing: AAR, MK and RJK; Visualization: AAR and RJK; Supervision: RJK; Project administration: RJK; Funding acquisition: RJK

## Competing interests

The authors declare no competing or financial interests

## Notes

### Competing Interest Statement

The authors have declared no competing interest.

### Summary of Updates

We have reanalyzed all the crystal cell differentiation data and plotted it as crystal cell differentiation index in all the main and supplementary figures for uniformity. We have revised our graphical summary/model. We have also revised the text to remove any extrapolations. The title, abstract and discussion section have also been revised.

