## Supplementary Information for "Gcn5 and mTORC1 mediated regulation of autophagy controls *Drosophila* blood cell homeostasis"

#### Supplemental Information

##### Supplementary methods:

###### ***Drosophila* Genetics**

All the *Drosophila* stocks and crosses were maintained at 25°C, in a standard cornmeal diet. *Canton S* was used as wild type control. The fly stocks used were *gcn5E333st/TM3* (BL-9333; RRID: BDSC\_9333), *gcn5C137Y/TM3* (BL-9335; RRID: BDSC\_9335), *UAS-Gcn5RNAi* (BL-9332; RRID: BDSC\_9332), *UAS-Gcn5FLAG*, *UAS-Gcn5ΔHAT*, *UAS-Gcn5ΔPcaf*, *UAS-Gcn5ΔBromo*, *UAS-Gcn5ΔAda* (Clement Carre, Sorbonne Universite, Paris), *UAS-TFEB RNAi* (Bhupendra Shravage, Agharkar Research Institute Pune, India), *UAS-Atg8aRNAi* (BL-34340; RRID:BDSC\_34340), *UAS-Atg5RNAi* (BL-34899; RRID:BDSC\_34899), *UAS-Atg18aRNAi* (BL-34714; RRID:BDSC\_34714), *tep4Gal4GFP*, *collierGal4mCD8GFP*, *DomeGal4mCD8GFP*, *HHLTGal4GFP*, *HmlGal4GFP*, *HmldeltaGal4GFP*, *LozengeGal4Mcd8gfp* (Lucas Waltzer, Université Clermont Auvergne, France), *UAS-TOR RNAi* (BL-33951 RRID:BDSC 33951), *UAS-raptor RNAi* (BL-41912 RRID:BDSC 41912), *UAS-rheb on II* (gift from Mohit Prasad). *UAS-Gcn5RNAi* or *UAS-Gcn5FLAG* were crossed with tissue-specific GAL4 lines to induce Gcn5 knockdown or over-expression.

###### **Generation of the E333st/E333st homozygous mutant:**

For the generation of the *E333st/E333st* homozygous mutant, the *E333st/TM3ser* heterozygous mutants were crossed with *tft/cyoGFP*; *UG3/TM3serGFP*. The F1 (*E333st/UG3*) were further screened for scorable marker i.e. non-serrate phenotype and inter crossed to generate *E333st/E333st* homozygotes.

###### **Antibodies**

Antibodies used were mouse anti-P1 (1:100, kind gift from Dr. Istvan Ando), mouse anti-Hindsight (1:25, 1G9 – DSHB; RRID:AB\_528278), mouse anti-Antp (1:25, 8C11- DSHB; RRID:AB\_528083), mouse anti-γ2AX (1:400, UNC93-5.2.1 - DSHB; RRID:AB\_2618077), mouse anti-DYKDDDDK Tag (anti-FLAG) (1:100, Thermo Fisher, RRID:AB\_1957945), anti-P62/SQSTM1 (1:250, Proteintech, RRID:AB\_10694431), anti-ATG8 (1:200, Sigma Aldrich, RRID:AB\_2939040), rabbit anti-dGcn5 (1:100, gift from Clement Carre) for immunofluorescence based experiments. Normal Goat Serum (HIMEDIA, RM10701) was used as the blocking agent. Alexa-Fluor 568 conjugated secondary antibodies – Goat anti- mouse 568 (1:400, Invitrogen, RRID:AB\_144696), and Goat anti-rabbit 568 (1:400, Invitrogen, RRID:AB\_10563566) were used for immunofluorescence based experiments. For Western blotting, antibody concentrations used were as follows: anti-dGcn5 (1:4000), anti-p62 (1:3000), anti-ATG8 (1:4000), and anti-β-actin (1:5000). Appropriate HRP conjugated secondary antibodies – Goat anti-Mouse (Invitrogen, RRID:AB\_228307) and Goat anti-Rabbit (Invitrogen, RRID:AB\_228341) were used at 1:5000 concentration.

###### **Lymph Gland Dissection and Immunohistochemistry**

Wandering third instar larvae were used for lymph gland dissections as discussed in (1). The dissections were performed in phosphate buffer saline (PBS), fixed in 4% paraformaldehyde, followed by washes with PBS containing 0.3% Triton-X (PBST). The samples were then blocked in 20% normal goat serum for 20 minutes at room temperature followed by overnight primary antibody incubation at 4°C. This was followed by PBST washes, blocking, and treatment with appropriate Alexa-Fluor conjugated secondary antibody incubation for two hours at room temperature. The LGs were then mounted in vectashield mounting medium containing DAPI (Vector Laboratories, RRID:AB\_2336790).

##### **Image acquisition and analysis of various Lymph Gland parameters**

Confocal images were captured using either Zeiss LSM 780, Leica SP8, or Nikon AX confocal microscope. Z projection of the confocal images were used for estimating various lymph gland parameters using ImageJ/Fiji software. Plasmatocyte Differentiation Index was estimated by measuring the percentile of P1 positive area divided by the total area of the primary lobe. The Prohemocyte Index was estimated by measuring the percentile of Tep4-GFP positive area divided by the total area of primary lobe. Freehand selection tool was used for measuring the area of the plasmacytocytes or the prohemocytes. For the quantification of Antp, Hnt, and γH2AX, the positive signals for respective markers were manually counted using the multi-point tool. For prohemocyte index quantitation, the GFP positive area marked by progenitors was selected, measured and it was further divided by the total area of the primary lobe. The crystal cell differentiation index was quantified by calculating the number of Hnt positive crystal cells/ total number of cells X 100. The LG quantifications were done for individual primary lymph gland lobes.

##### **Quantification of p62 and Atg8:**

Three ROIs were selected from larval LGs stained with p62 or Atg8. 5 cells per ROI were used to quantify the p62 and Atg8 positive puncta, further an average of the number of puncta per cell was quantified per ROI as the total number of positive puncta/total number of cells. These data points were further compared with the control to determine statistical significance.

##### **Protein extraction and Western Blotting**

Around 10-12 whole larvae were lysed in RIPA buffer containing protease inhibitors followed by homogenization and sonication (30s-ON, 30s-OFF x 10 cycles). The lysate was then centrifuged at 10000rpm for 5 minutes at 4°C. The supernatant was collected and quantified using BCA Protein Assay kit (Thermo Scientific, 23227) and stored at -80°C. 50μg of proteins were loaded and separated on SDS-PAGE and transferred onto a polyvinylidene difluoride (PVDF) membrane, blocked in 5% BSA for 1 hour at room temperature. The membranes were then probed with different primary antibodies. Blots were developed using an Enhanced Chemiluminescent substrate (Thermo Scientific, 34580) in Chemidoc (Bio-Rad). All the Western blotting experiments were done in biological triplicates.

##### **Quantitative Real-Time PCR**

RNA was extracted from 500 adult flies, by removing the head and perfusing the flies in cold PBS as described previously (3). The hemolymph was pelleted by centrifuging at 1500rpm for 5 minutes at 4°C. The supernatant was removed and the hemolymph pellet was lysed in TRIzol (Ambion – life technologies, 11596018). The lysates were stored at -80°C. Batches of 100 flies or more were done and once the hemolymph from all 500 flies were done, RNA was isolated by pooling all the aqueous layers post-chloroform treatment, followed by RNA isolation according to the manufacturer's protocol. RNA yield was quantified using Nanodrop. 1μg of mRNA was reverse transcribed using oligo-dT primers (Promega, C110A) and ImProm-II (Promega, A3800). Quantitation of the transcripts was done using SYBR green chemistry in Quantstudio 5 RT PCR system (Thermo Fischer Scientific). The data was analyzed using the $\Delta\Delta$Ct method and relative mRNA expression was normalized to rp49. Fold change calculations were done in comparison to wildtype control. The experiment was done in biological triplicates.

##### **Chemical treatment**

For drug treatments, early third instar larvae were collected and transferred to vials containing distilled water and starved for 2 hours. The larvae were then transferred to food containing corresponding drugs to be used for treatment. Chemicals used include Rapamycin (40μg/ml,

R0395, Sigma-Aldrich) dissolved in absolute ethanol, 3BDO (200μM, SML1687, Sigma Aldrich) dissolved in DMSO, and Chloroquine diphosphate salt (2.5mg/ml, C6628, Sigma-Aldrich) dissolved in distilled water. For control media, post starvation, the late second instar larvae were fed on food containing respective diluents alone. For starvation experiment, the larvae were starved for 2 hours in vials containing distilled water they were then transferred to a vial containing 1ml of media mixed with chloroquine. The larvae were treated for 16 hours and were further dissected for immunostaining and then lysate was made for western blotting. For the high-fat diet experiments, the late first instar larvae were directly transferred to and reared in food containing 20% weight per volume of food-grade coconut oil. These larvae fed on a high fat diet were used for further experiments. For each of the drug treatment experiments, atleast 12 larvae were used for analysis and the treatment was done for 16 hours.

##### Statistical Analysis

Immunofluorescence based experiments and their analysis was performed on atleast 10 lymph glands dissected from wandering third-instar larvae. For Western Blotting and qPCR-based analysis, the experiments were done in triplicates and then analyzed. Statistical analysis was performed using the GraphPad Prism Version 9 software (RRID:SCR\_002798). For analysis of statistical significance each experimental sample was tested with its respective control in a given experimental setup for all the data in each of the figures in order to estimate the P value. P values were determined by using a two-tailed unpaired Student's t-test with Welch's correction. \*\*\*\* indicates $p < 0.0001$, \*\*\* indicates $p < 0.001$, \*\* indicates $p < 0.01$, \* indicates $p < 0.05$, ns (non-significant) indicates $p > 0.05$. Mutant genotypes were compared to the wild- type controls and the knockdown or overexpression genotypes were compared to their respective parental Gal4 controls that were crossed to wild-type for all the statistical analysis performed. The experiments where chemical treatment has been given have been compared to the respective controls. No statistical method was used to predetermine the sample size and the experiments were not randomized. The sample size for each of the experiments has been indicated in the respective figure legends.

##### List of primers:

qRT primers in 5' to 3' direction

|  |  |
| --- | --- |
| Atg1 Forward | GGCAGTGGATCGGAGAACAA |
| Atg1 Reverse | CATCAGCGTCTCTTCGGACA |
| Atg4a Forward | TGGTTCTGTTGAAGCTGACC |
| Atg4a Reverse | CCTCATTGGGTGCTCCACTT |
| Atg4b Forward | TTGCACGAGTAAGTTCAAGCA |
| Atg4b Reverse | CAAGGCACATGGGGTTTTGG |

|  |  |  |
| --- | --- | --- |
| Atg5 | Forward | AGGGAAAAGGTGCCAGTCAG |
| Atg5 | Reverse | GATTTTTCGGTTCGGCTCGG |
| Atg7 | Forward | CCGAAGGCGTCATCTTTGTT |
| Atg7 | Reverse | GTTGGATACTCCGGGTCGTG |
| Atg9 | Forward | TCAGTACCAGCAGAAGCACG |
| Atg9 | Reverse | GCAGTGCATCACAAAGGCAA |
| Atg10 | Forward | GGCTTGTTTCCCACGCAAAA |
| Atg10 | Reverse | TCCGCACTGCGAGATACCTTT |
| Atg13 | Forward | GGCGTTGCACAAATGACAGT |
| Atg13 | Reverse | CGTTCCGCTGCATTTAGACG |
| Atg14 | Forward | ACTGATGCTGATGCCTTTCCA |
| Atg14 | Reverse | GCGATGGTAGACTGCTGGTT |
| Atg10 | Forward | TAAATGCCGAGGTGGCAAGG |
| Atg10 | Reverse | TCTGTGTGCCTGGAAGAACAA |
| Atg18a | Forward | GCACGCCAAGACCATGATTC |
| Atg18a | Reverse | TTTAGTCCCCGTCGCAGTTC |

#### SI figure legends:

##### Figure S1. Blood cell homeostasis is affected in the whole animal *gcn5* heterozygous mutants.

PSC cell numbers marked by Antennapedia (red) in *gcn5*[C137Y/+]<sup>1</sup> (B-B') or *gcn5*[E333st/+]<sup>2</sup> (C-C') as compared to *wildtype* (A-A'). Graphical representation of PSC cell numbers of the *gcn5* mutants compared to the *wildtype* (D), n=24 for *wildtype*, n=33 for *gcn5*[C137Y/+]<sup>1</sup>, and n=41 for *gcn5*[E333st/+]<sup>2</sup> for PSC cell numbers quantification. Plasmatocyte differentiation was marked by P1 (red) in *gcn5*[C137Y/+]<sup>1</sup> (F-F') or *gcn5*[E333st/+]<sup>2</sup> (G-G') compared to the *wildtype* (E-E'). Graphical representation of plasmatocyte differentiation index of the *gcn5* heterozygous mutants compared to *wildtype* (H), n=24 for *wildtype*, n=20 for *gcn5*[C137Y/+]<sup>1</sup>, and n=24 for *gcn5*[E333st/+]<sup>2</sup> for quantification. Crystal cell differentiation marked by Hnt (red) in *gcn5*[C137Y/+]<sup>1</sup> (J-J') or *gcn5*[E333st/+]<sup>2</sup> (K-K') compared to *wildtype* (I-I'). Graphical representation of Crystal cell differentiation index for *gcn5* heterozygous mutants compared to *wildtype* (L), n=22 for *wildtype*, n=28 for *gcn5*[C137Y/+]<sup>1</sup>, and n=23 for *gcn5*[E333st/+]<sup>2</sup> for quantification. Cells undergoing DNA damage marked by γH2AX (red) in *gcn5*[C137Y/+]<sup>1</sup> (N-N') or *gcn5*[E333st/+]<sup>2</sup> (O-O') as compared to *wildtype* (M-M'). Graphical representation of γH2AX marked DNA damage of the heterozygous *gcn5* mutants compared to the *wildtype* (P), n=31 for *wildtype*, n=42 for *gcn5*[C137Y/+]<sup>1</sup>, and n=28 for *gcn5*[E333st/+]<sup>2</sup> for quantification. Nuclei are stained with DAPI (Blue). n represents the number of individual primary lobes of the LG. Individual data points in the graphs represent individual primary lobes of the LG. Values are mean ± SD, and asterisk marks statistically significant differences (\*p<0.05; \*\*p<0.01; \*\*\*p<0.0001, Student's t-test with Welch's correction). Scale Bar: 50μm (A-O').

##### Figure S2. Validation of Gcn5 knockdown and over-expression constructs

Gcn5 (red) expression upon Hml-Gal4 mediated Gcn5 knockdown (B-B') or over-expression (C-C') in the hml population, compared to *wildtype* (A-A'). FLAG (red) expression upon Gcn5 over-expression (E-E') in the hml population, compared to *wildtype* (D-D'). Nuclei are stained with DAPI (Blue). Scale Bar: 50μm (A-E').

##### Figure S3. Collier mediated modulation of Gcn5 levels in the PSC alters LG homeostasis.

PSC cell population marked by Antennapedia (red) upon knockdown (B-B') or over-expression (C-C') of Gcn5 in the Collier population (green) compared to *wildtype* (A-A'). Plasmatocyte differentiation marked by P1 (red) upon knockdown (E-E') or over-expression (F-F') of Gcn5 in the Collier population (green) compared to *wildtype* (D-D'). Crystal cell differentiation marked by Hnt (red) upon knockdown (H-H') or over-expression (I-I') of Gcn5 in the Collier population (green) compared to *wildtype* (G-G'). Cells undergoing DNA damage marked by γH2AX (red) upon knockdown (K-K') or over-expression (L-L') of Gcn5 in the Collier population (green) compared to *wildtype* (J-J'). Graphical representation of PSC cell numbers upon modulation of Gcn5 levels in the Collier population compared to *wildtype* (M), n=21 for *wildtype*, n=29 for Gcn5 knockdown, and n=22 for Gcn5 over-expression. Graphical representation of plasmatocyte differentiation Index upon modulation of Gcn5 levels in Collier population compared to *wildtype* (N), n=23 for *wildtype*, n=23 for Gcn5 knockdown, and n=20 for Gcn5 over-expression. Graphical representation of Crystal cell differentiation index upon modulation of Gcn5 levels in the Collier population compared to *wildtype* (O), n=34 for *wildtype*, n=24 for Gcn5 knockdown, and n=24 for Gcn5 over-expression. Graphical representation of γH2AX positive cells upon modulation of Gcn5 levels in the Collier population compared to *wildtype* (P), n=29 for *wildtype*, n=27 for Gcn5 knockdown, and n=21 for Gcn5 over-expression. Nuclei are stained with DAPI (blue). PSC cell specific Gcn5

modulation was performed using Collier-Gal4. n represent the number of individual primary lobes of the LG. Individual data points in the graphs represent individual primary lobes of the LG. Values are mean ± SD, and asterisk marks statistically significant differences (\*$p$<0.05; \*\*$p$<0.01; \*\*\*$p$<0.001, Student's t-test with Welch's correction). Scale Bar: 50μm (A-L')

**Figure S4. Hml $\Delta$ mediated modulation of Gcn5 levels in the differentiated blood cell population results in altered hematopoiesis.**

PSC cell population marked by Antennapedia (red) upon knockdown (B-B') or over-expression (C-C') of Gcn5 in the HmlΔ population (green) compared to wildtype (A-A'). Plasmatocyte differentiation marked by P1 (red) upon knockdown (E-E') or over-expression (F-F') of Gcn5 in the HmlΔ population (green) compared to wildtype (D-D'). Crystal cell differentiation marked by Hnt (red) upon knockdown (H-H') or over-expression (I-I') of Gcn5 in the HmlΔ population (green) compared to wildtype (G-G'). Cells undergoing DNA damage marked by γH2AX (red) upon knockdown (K-K') or over-expression (L-L') of Gcn5 in the HmlΔ population (green) compared to wildtype (J-J'). Graphical representation of PSC cell numbers upon modulation of Gcn5 levels in the HmlΔ population compared to wildtype (M), n=24 for wildtype, n=25 for Gcn5 knockdown, and n=36 for Gcn5 over-expression. Graphical representation of plasmatocyte differentiation Index upon modulation of Gcn5 levels in HmlΔ population compared to wildtype (N), n=20 for wildtype, n=20 for Gcn5 knockdown, and n=21 for Gcn5 over-expression. Graphical representation of Crystal cell differentiation index upon modulation of Gcn5 levels in the HmlΔ population compared to wildtype (O), n=20 for wildtype, n=20 for Gcn5 knockdown, and n=23 for Gcn5 over-expression. Graphical representation of γH2AX positive cells upon modulation of Gcn5 levels in the HmlΔ population compared to wildtype (P), n=23 for wildtype, n=29 for Gcn5 knockdown, and n=27 for Gcn5 over-expression. Nuclei are stained with DAPI (blue). HmlΔ-Gal4 was used for differentiated blood cells specific Gcn5 modulation. n represent the number of individual primary lobes of the LG. Individual data points in the graphs represent individual primary lobes of the LG. Values are mean ± SD, and asterisk marks statistically significant differences (\*$p$<0.05; \*\*\*$p$<0.001; \*\*\*\*$p$<0.0001, Student's t-test with Welch's correction). Scale Bar: 50μm (A-L').

**Figure S5: GCN5 positively regulates crystal cell differentiation cell-autonomously**

GCN5 (magenta) expression in lz-GFP (Green) positive crystal cells in larval LGs where lz-Gal4 drives UAS-GFP in wild type genetic background (A-B). Crystal cell differentiation marked by Hindsight (Hnt, magenta) upon GCN5 over-expression using lz- Gal4GFP (D) as compared to control (C) and quantitated as crystal cell differentiation index per LG lobe represented by n, where n=31 for wildtype and n=36 for GCN5 overexpression. (G). GCN5 (magenta) expression in posterior LG lobes marked by GFP (Green) driven by tep4- Gal4 (E-F). Nuclei = DAPI (Blue) and GFP driven by either lz-Gal4 or tep4-Gal4. Values are mean ± SD, and asterisk marks statistically significant differences with \*\*\* as $p$<0.001 analyzed by Student's t- test with Welch's correction. Scale Bar: 30μm (A-B), 50 μm (C-F).

**Figure S6. Prohemocyte-specific expression of Gcn5 domain deletion constructs leads to increased DNA damage.**

Cells undergoing DNA damage marked by γH2AX (red) upon expression of different Gcn5 domain deletion constructs in the tep-GFP population using tep4-Gal4 (green) (B-E') as compared to wildtype (A, A'). Graphical representation of Cells undergoing DNA damage marked by γH2AX (F), where n=27 for wildtype, n=26 for Gcn5ΔHAT expression, n=27 for Gcn5ΔPcaf expression, n=23 for Gcn5ΔBromo expression, and n=25 for Gcn5ΔAda expression in the tep population. Nuclei are stained with DAPI (blue). n represents the number of individual primary lobes of the LG. Individual data points in the graphs represent individual primary lobes of the LG. Values are mean ± SD, and asterisk marks statistically significant differences (\*\*$p < 0.01$; \*\*\*$p < 0.001$; \*\*\*\*$p < 0.0001$, Student's t-test with Welch's correction). Scale Bar: 50μm (A-E')

**Figure S7. Gcn5 over-expression leads to suppression of autophagy-related genes**
Transcript levels of different autophagy effector genes in Hml-Gal4 mediated Gcn5 over-expression genetic background as compared to the Hml-Gal4 X wt control. rp49 was used as an endogenous control to normalize the mRNA expression.

**Figure S8: Prohemocyte- specific genetic abrogation of autophagy affects the niche cell numbers and induces DNA damage.**

PSC/niche cells marked by Antennapedia (red) upon tep4-Gal4 mediated knockdown of TFEB (C-C'), Atg8a (E-E'), Atg5 (G-G'), or Atg18 (I-I') in the tep population (green) as compared to wildtype (A-A'). DNA damage marked by Gamma-H2Ax (red) upon tep4-Gal4 mediated knockdown of TFEB (D-D'), Atg8a (F-F'), Atg5 (H-H'), or Atg18 (J-J') in the tep population (green) as compared to wildtype (B-B'). Graphical representation of niche cell numbers upon knockdown of TFEB (n=40), Atg8a (n=36), Atg5 (n=39), Atg18 (n=41) in the tep population, compared to the wildtype (n=33) (K). Graphical representation of Gamma-H2Ax positive foci upon knockdown of TFEB (n=37), Atg8a (n=34), Atg5 (n=33), or Atg18 (n=36) in the tep population, compared to the wildtype (n=35) (L). Graphical representation of prohemocyte index upon knockdown of TFEB (n=30), Atg8a (n=30), Atg5 (n=30), Atg18 (n=30) in the tep population, compared to the wildtype (n=33) (M). Nuclei are stained with DAPI (blue). n represents the number of individual primary lobes of the LG. Individual data points in the graphs represent individual primary lobes of the LG. Values are mean ± SD, and asterisk marks statistically significant differences (\*$p < 0.05$; \*\*\*$p < 0.001$; \*\*\*\*$p < 0.0001$, Student's t- test with Welch's correction). Scale Bar: 50μm (A-J')

**Figure S9: Autophagy levels in the LG hemocytes are affected upon systemic Chloroquine treatment**

Chloroquine treated tep4-Gal4GFP X wt larvae expressing GFP (Green) driven by tep4-Gal4 showing p62 or Atg8 positive puncta (magenta) in LG hemocytes (B-B', D-D') as compared to vehicle treated (A-A', C-C') and represented as number of p62 or Atg8 positive puncta per cell (n), where n= 255 cells for vehicle treated and n=300 cells for chloroquine treated p62 positive cells and n= 360 cells for vehicle treated and n= 390 cells for chloroquine treated Atg8 positive cells respectively.(E and F). Nuclei = DAPI (Blue) and GFP driven by tep4-Gal4. Values are mean ± SD, and asterisk marks statistically significant differences with \*\*\*\*$p < 0.0001$ analyzed by Student's t- test with Welch's correction. Scale Bar: 30μm (A-D')

**Figure S10: Genetic modulation of mTORC1 activity in the hematopoietic progenitors regulates blood cell differentiation**

Plasmatocyte differentiation marked by P1 (magenta) upon tep4-Gal4 mediated depletion of tor (B) or raptor (C) or over-expression of Rheb (D) as compared to the wild type control (A) quantitated as plasmatocyte differentiation index per LG lobe where n represents the number of primary lobes of LG, n= 45 for wildtype, n= 39 for tor knockdown, n= 43 for raptor knockdown and n=44 for Rheb overexpression. (I). Crystal cell differentiation marked by Hindsight (Hnt, magenta) upon tep4-Gal4 mediated depletion of tor (F) or raptor (G) or over-expression of Rheb (H) as compared to the wild type control (E) quantitated as crystal cell differentiation index per LG lobe(n), where n represents the number of primary lobes of LG, n= 31 for wildtype, n= 42 for tor knockdown, n=30 for raptor knockdown, n=28 for Rheb overexpression. (J). Nuclei = DAPI (Blue) and GFP is driven by tep4-Gal4. n = number of individual primary lobes of the LG. Individual data points in the graphs represent individual primary lobes of the LG. Values are mean ± SD, and asterisk marks statistically significant differences (\*$p < 0.05$; \*\*\*$p < 0.001$; \*\*\*\*$p < 0.0001$, ns = non-significant. Student's t- test with Welch's correction). Scale Bar: 50μm (A-H)

**Figure S11: Genetic modulation of mTORC1 activity in the hematopoietic progenitors alters niche cell numbers and causes DNA damage**

PSC/niche cell numbers marked by Antennapedia (Antp, magenta) upon tep4-Gal4 mediated depletion of tor (B) or raptor (C) or over-expression of Rheb (D) as compared to the wild type control (A) quantitated as Antp positive niche cell numbers per LG lobe represented by n, where n= 38 for wildtype, n= 37 for tor knockdown, n=42 for raptor knockdown, n=37 for Rheb overexpression. (I). γ-H2Ax foci positive cells showing DNA damage (magenta) upon tep4-Gal4 mediated depletion of tor (F) or raptor (G) or over-expression of Rheb (H) as compared to the wild type control (E) quantitated as total number of γ-H2Ax positive foci per LG lobe represented by n, where n= 40 for wildtype, n= 43 for tor knockdown, n=34 for raptor knockdown, n=37 for Rheb overexpression. (J). Nuclei = DAPI (Blue) and GFP is driven by tep4-Gal4. n = number of individual primary lobes of the LG. Individual data points in the graphs represent individual primary lobes of the LG. Values are mean ± SD, and asterisk marks statistically significant differences (\*$p < 0.05$; \*\*\*$p < 0.001$; \*\*\*\*$p < 0.0001$, ns = non-significant. Student's t- test with Welch's correction). Scale Bar: 50μm (A-H)

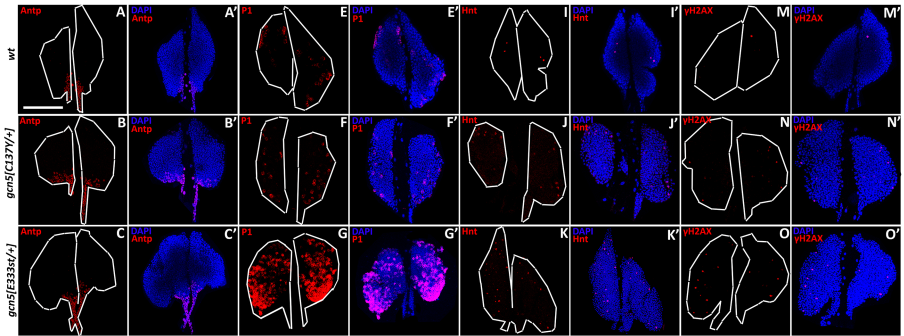

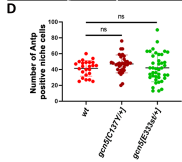

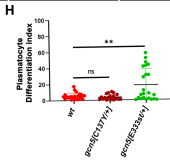

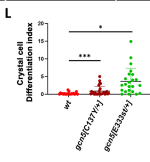

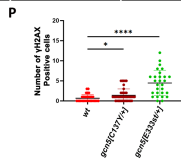

*HmlGal4 x wt*

*HmlGal4>UAS-Gcn5RNAi*

*HmlGal4>UAS-Gcn5FLAG*

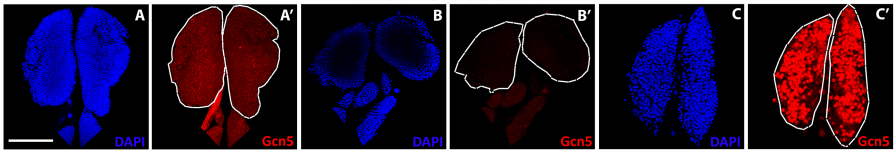

*HmlGal4 x wt*

*HmlGal4>UAS-Gcn5FLAG*

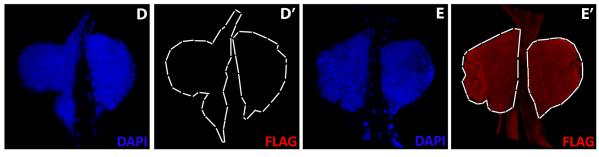

### CollierGal4GFP x wt

### CollierGal4GFP x UAS-Gcn5RNAi

### CollierGal4GFP x UAS-Gcn5

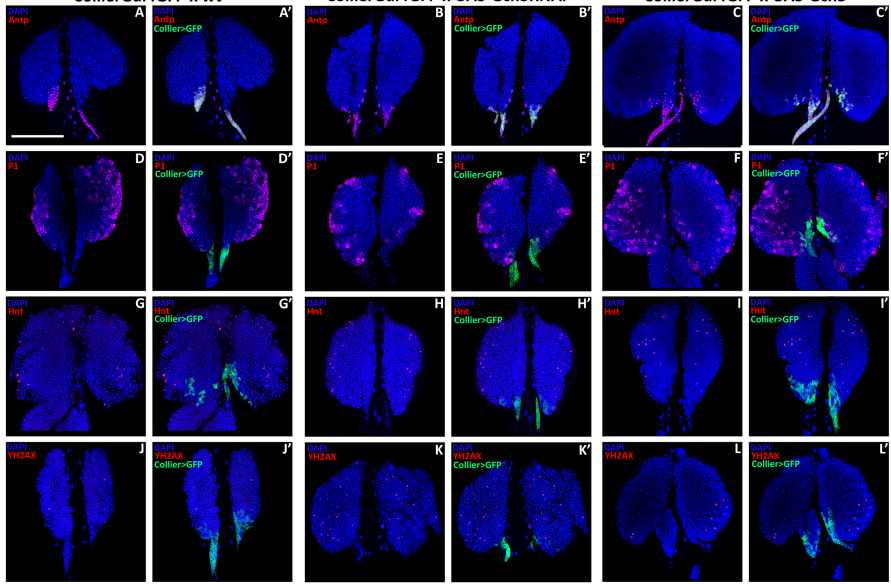

M

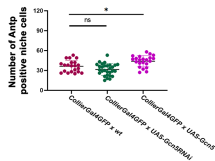

N

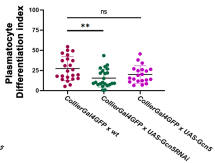

O

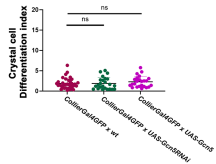

P

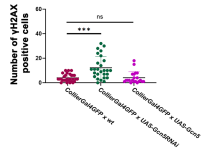

### HmldeltaGal4GFP x wt

### HmldeltaGal4GFP x UAS-Gcn5RNAi

### HmldeltaGal4GFP x UAS-Gcn5

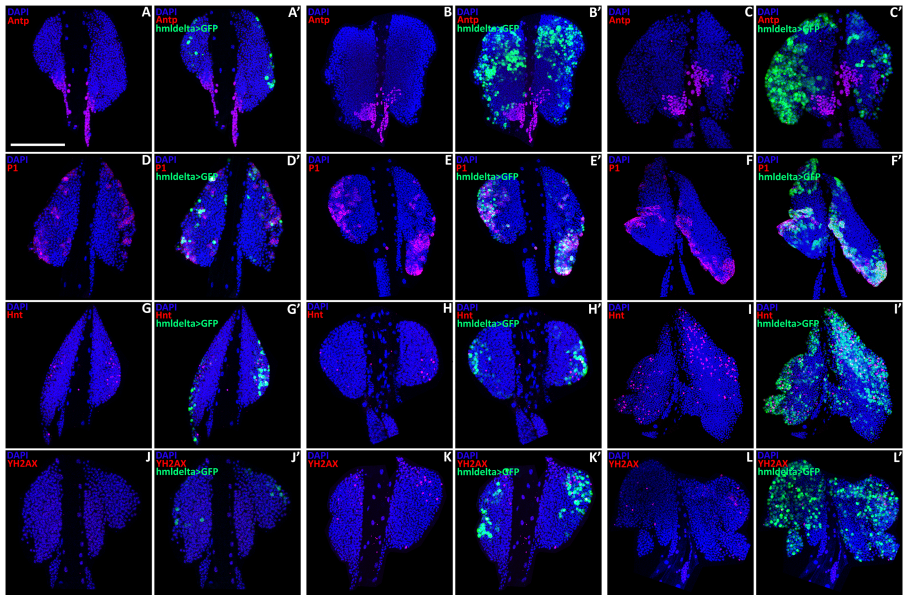

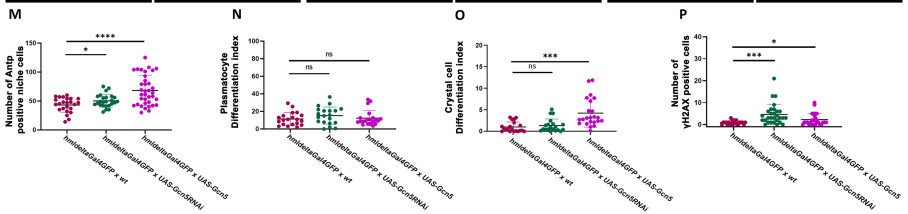

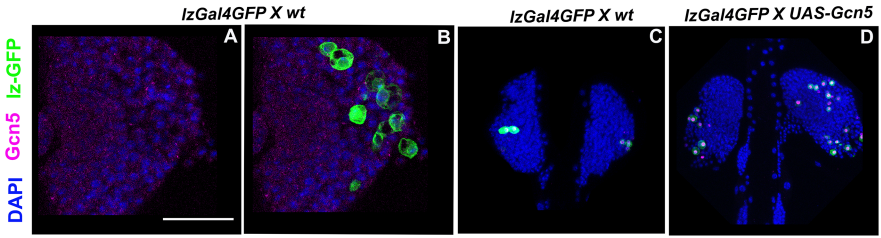

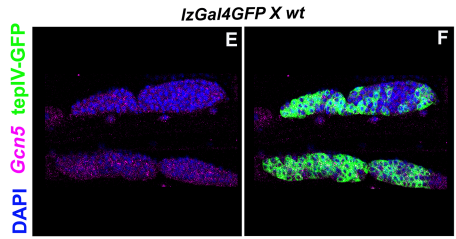

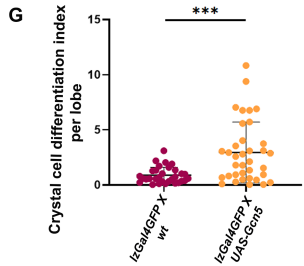

*Tep4Gal4GFP x wt*

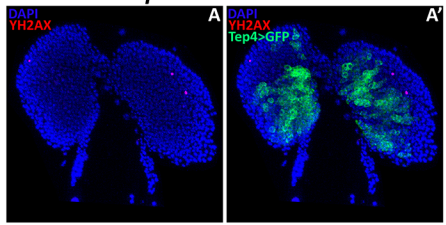

*Tep4Gal4GFP x UAS-Gcn5ΔHAT*

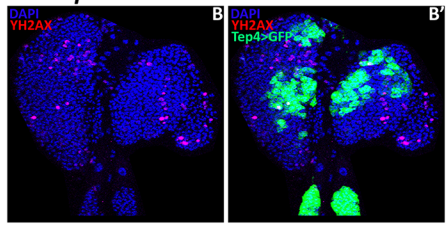

*Tep4Gal4GFP x UAS-Gcn5ΔPcaf*

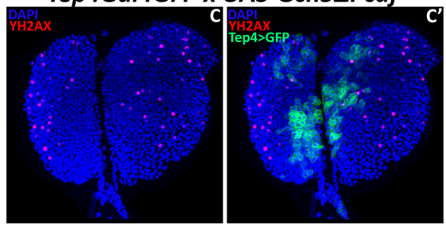

*Tep4Gal4GFP x UAS-Gcn5ΔBromo*

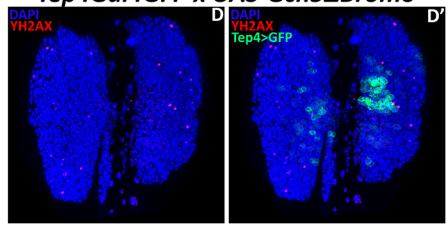

*Tep4Gal4GFP x UAS-Gcn5ΔAda*

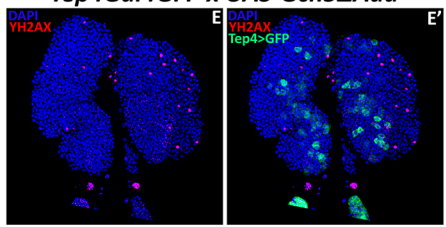

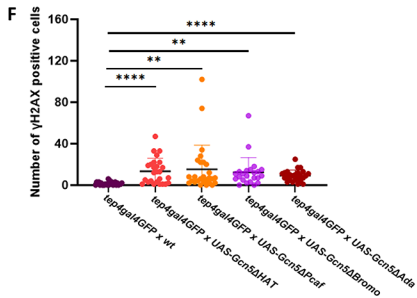

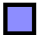

*HmlGal4>UAS-Gcn5*

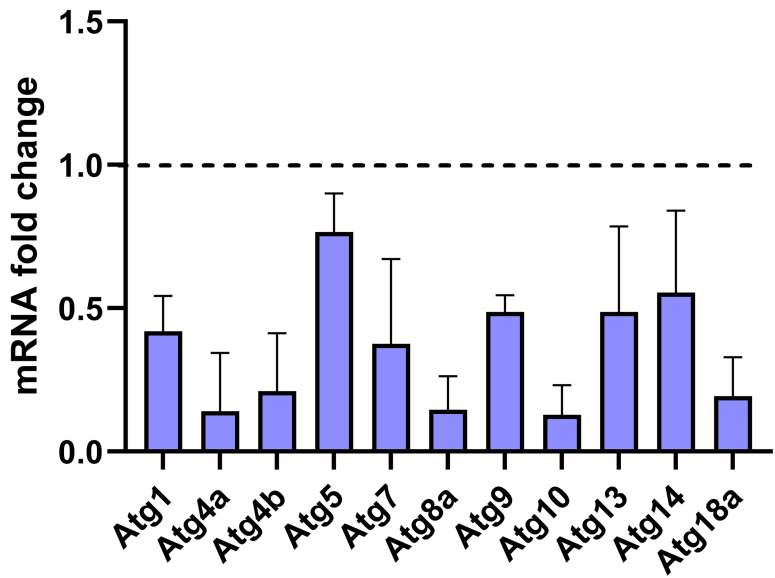

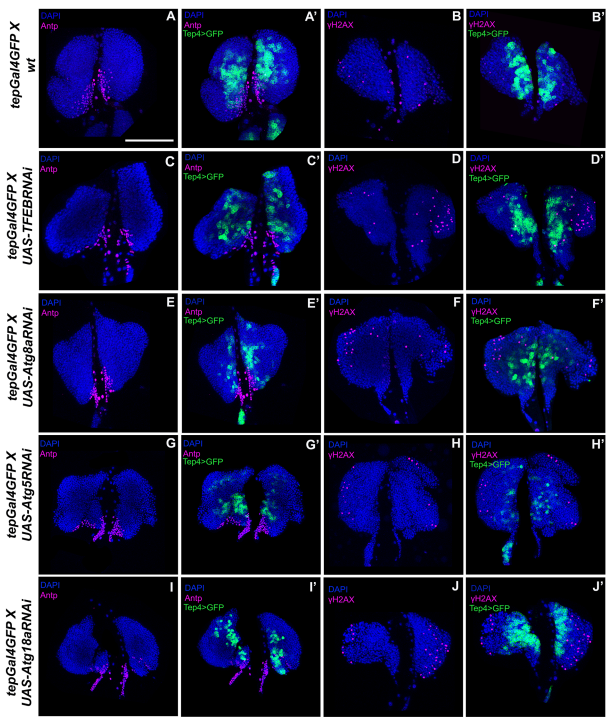

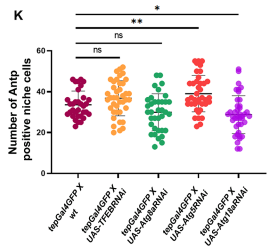

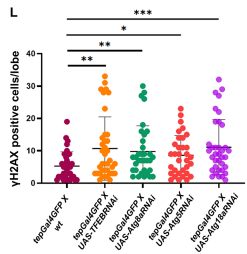

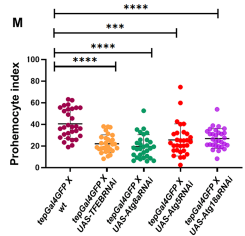

DAPI Atg8 tepIV-GFP DAPI p62 tepIV-GFP

DAPI Atg8 tepIV-GFP

DAPI Atg8 tepIV-GFP DAPI p62 tepIV-GFP

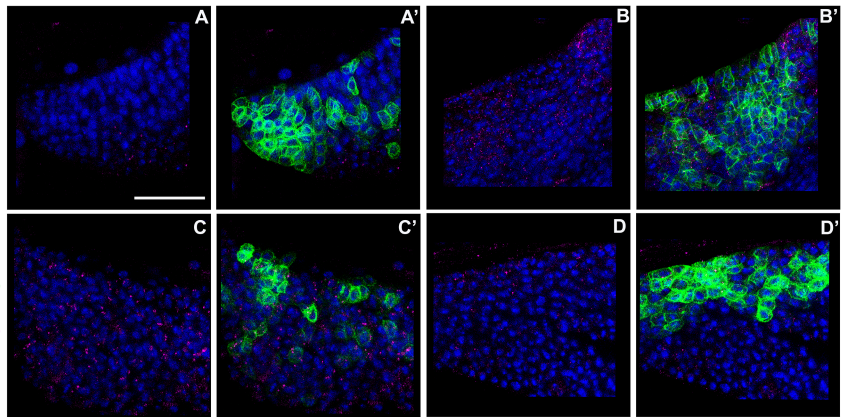

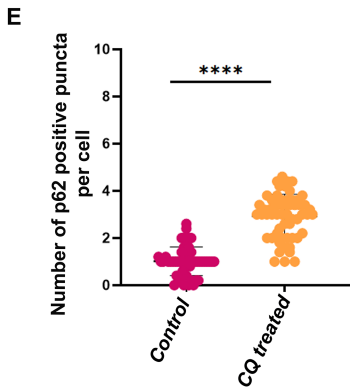

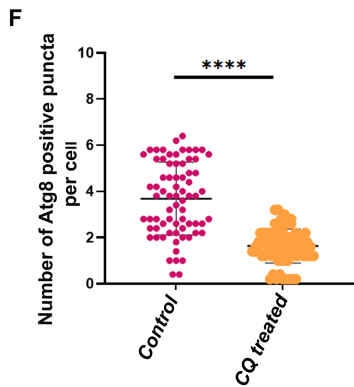

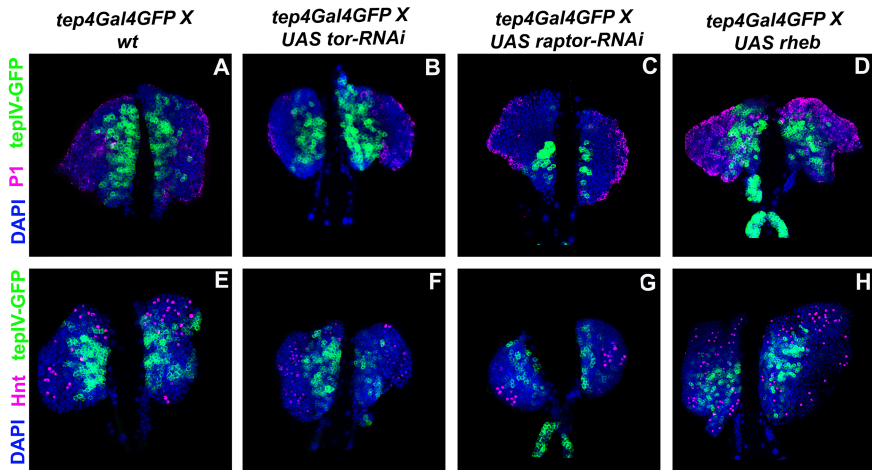

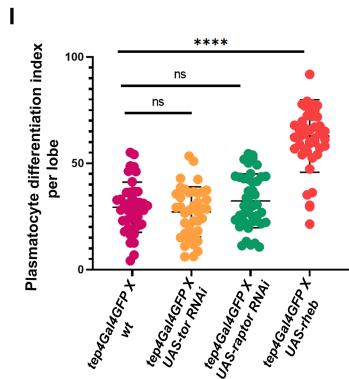

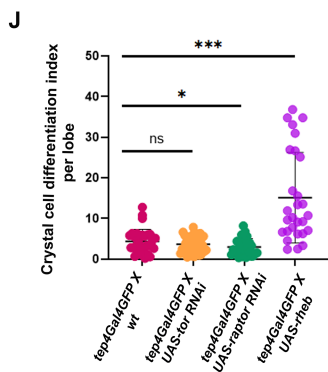

DAPI *yH2AX* *tepIV-GFP* Antp *tepIV-GFP*

*tep4Gal4GFP X*
*wt*

*tep4Gal4GFP X*
*UAS tor-RNAi*

*tep4Gal4GFP X*
*UAS raptor-RNAi*

*tep4Gal4GFP X*
*UAS rheb*

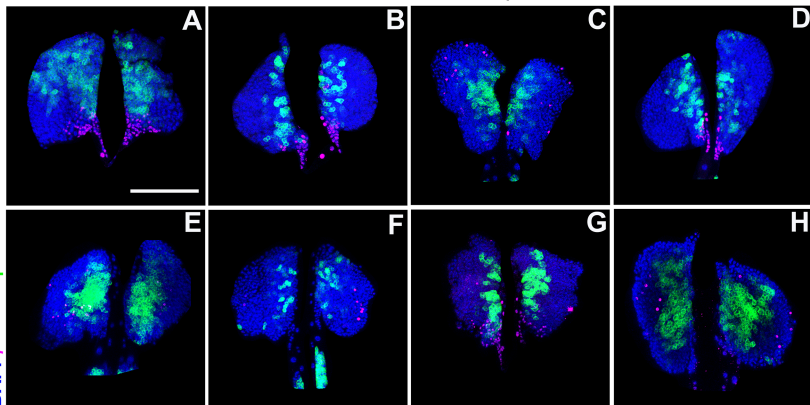

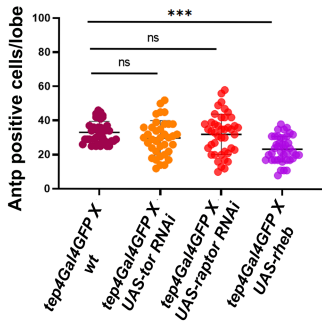

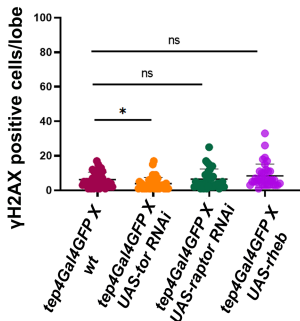
